# Sex differences define the molecular and cellular phenotypes of pain resolution in dorsal root ganglia

**DOI:** 10.64898/2025.12.01.691610

**Authors:** Felicitas Schlott, Beate Hartmannsberger, Thorsten Bischler, Tom Gräfenhan, Alexander Brack, Heike L. Rittner, Annemarie Sodmann, Robert Blum

**Affiliations:** Department of Neurology, University Hospital Würzburg, Würzburg, Germany; Department of Anesthesiology, Intensive Care, Emergency Medicine and Pain Therapy, Centre for Interdisciplinary Pain Medicine, University Hospital Würzburg, Würzburg, Germany; Core Unit Systems Medicine, University of Würzburg, Würzburg, Germany

**Keywords:** dorsal root ganglia, neuropathic pain, macrophages, satellite glial cells, sensory neurons, sex differences, bioimage analysis, deep learning, transcriptome analysis

## Abstract

The dorsal root ganglion (DRG), a key site for the initiation and maintenance of neuropathic pain, was examined for sex-dependent phenotypes in sensory neurons, satellite glial cells (SGCs), and local macrophages following traumatic nerve injury and during natural pain resolution. Systematic analysis of 7,495 DRG immunofluorescence images and 62 transcriptomes revealed pronounced sex-specific, multicellular DRG phenotypes, especially during pain resolution. System parameters, including tissue size and neuron density also showed sex-dependent differences. Neuropathic pain resolved without tissue or sensory neuron loss. After injury, macrophages invaded the space between sensory neurons and satellite glial cells (SGCs); this was partially reversed during pain resolution, particularly in males. In females, immune-related gene expression and macrophage phenotypes persisted longer, while SGC activation and contact to sensory neurons was more persistent in males. During resolution, synaptic and excitability-related processes were pronounced in both sexes. However, while injury responses were largely shared between sexes, the resolution phase displayed distinctly sex-specific molecular and cellular signatures.

**GRAPHICAL ABSTRACT:** 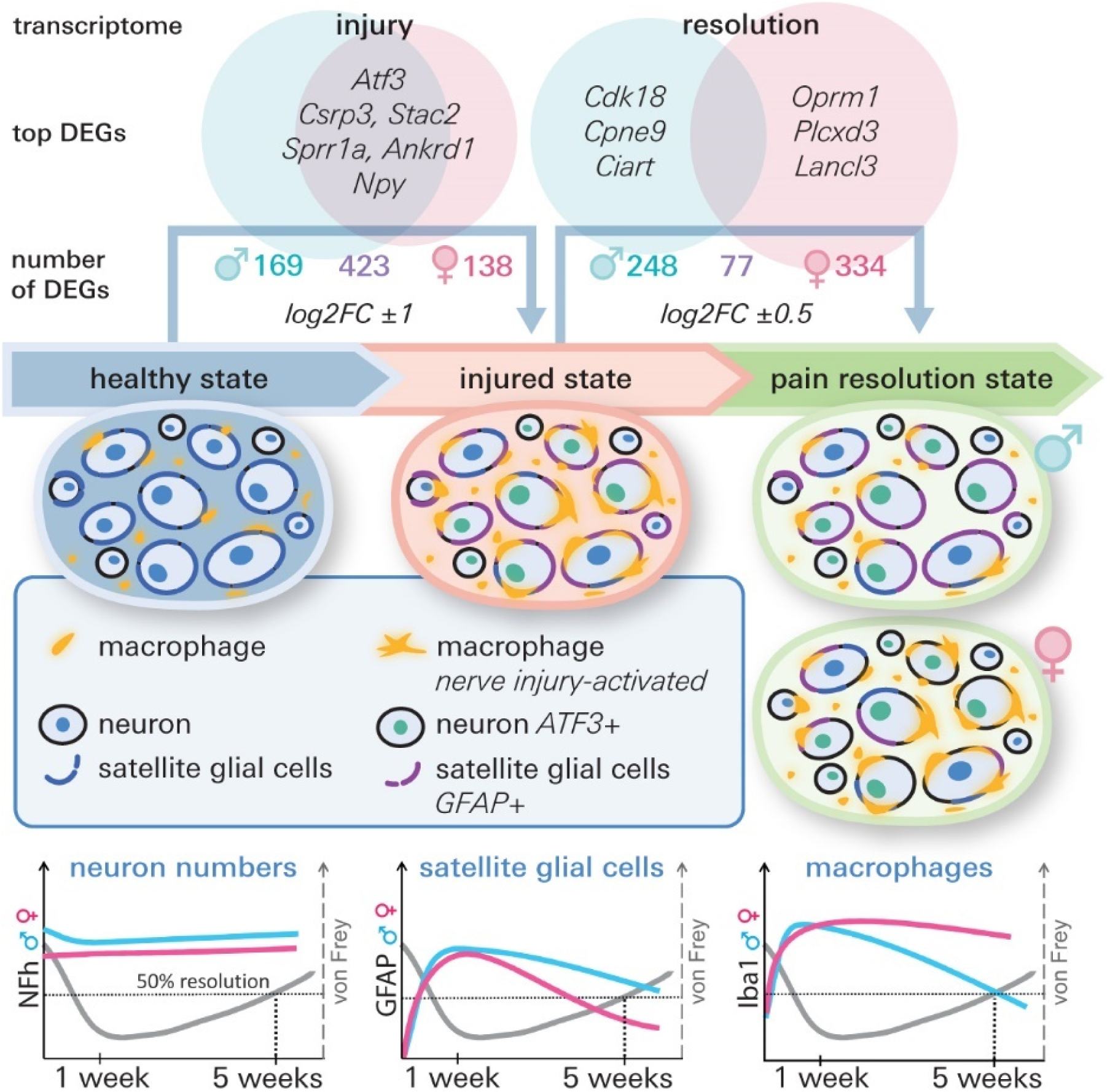

**In brief:** Analysis of ∼7,500 bioimages and 62 transcriptomes reveals pronounced sex differences in rat dorsal root ganglia during pain resolution after peripheral nerve injury.

**Highlights:**

- In both female and male rats, peripheral nerve injury and subsequent pain resolution occur in the dorsal root ganglia (DRG) without neuronal or tissue loss.
- Sex influences DRG tissue size, neuron density, immune and glial phenotypes, and molecular-cellular responses to nerve injury and pain resolution.
- Following injury, macrophages infiltrate the space between sensory neurons and satellite glial cells (SGCs); this process reverses during pain resolution, particularly in males.
- In females, immune phenotypes remain more stable throughout pain resolution, while SGC contact is reduced.
- Pain resolution involves not only the reversal of injury-induced cell changes but also the activation of resolution-specific gene programs related to synaptic signaling, neuronal excitation, and cell–cell communication.
- Sex differences on the molecular-cellular level are less prevalent after nerve injury but become prominent during pain resolution.

## INTRODUCTION

Neuropathic pain arises from lesions or diseases affecting the somatosensory nervous system (1). Currently, effective treatment options remain insufficient (2), allowing pain to frequently transit from acute to chronic states. Affecting up to ∼7% to 10% of the general population (3), chronic neuropathic pain imposes significant personal and socio-economic burden (4). Although chronic pain following nerve injury can improve naturally (5), the reasons why neuropathic pain resolves in some individuals but not in others remain unclear.

Pain resolution is a biological process encompassing both the onset of the pain and recovery to a state of low or absent pain. Experimental studies have identified several mechanisms that may facilitate pain resolution, including the active resolution of inflammation (6), or suggest potential restoration of sensory pathways or transcriptional profiles within sensory neurons (7–9). Mechanisms beyond inflammation resolution have also been proposed. For instance, restoring neuronal energy metabolism through mitochondrial transfer from macrophages to sensory neurons has been shown to contribute to natural pain resolution (10).

In humans, neuronal and non-neuronal cells in the dorsal root ganglion (DRG) show diverse responses to nerve injuries and pain. In painful diabetic neuropathy, significant sensory neuron loss occurs, and immune phenotypes might contribute to persistence of pain (11, 12). Similarly, in brachial plexus injuries, half of the patients exhibited complete DRG loss, while the rest retained sensory neurons, satellite glial cells, and macrophages (13). However, even after partial nerve reconstruction after injury, neither patient group showed pain resolution (13).

In animal neuropathic pain models, neuronal loss was observed after nerve injury (14, 15). For instance, in mice, painful nerve injuries were accompanied by DRG shrinkage and sensory neuron loss, particularly of non-peptidergic nociceptors (14). In rats, within 14 days post spared nerve injury (SNI), we observed no neuron loss (16) albeit pain does not resolve naturally after SNI (7).

An important but often underexplored biological variable of nociception, pain perception, and pain resolution is sex (17–20). Many pain syndromes are more prevalent in women (21, 22) and substantial evidence supports the existence of sex differences in pain experiences and in the transition from acute to chronic pain (17, 19, 23, 24). These differences may be linked to conserved biological mechanisms such as nociceptor transcriptomic signatures, sex hormone levels (25, 26), sexually dimorphic responses to CGRP (27), or sex-specific immune cell contributions by microglia (28) and T-cells (29). Notably, even life span under neuropathic pain conditions varies by sex (30).

In this study, we focused on sex-specific molecular and cellular phenotypes of sensory neurons, satellite glial cells and local macrophages in the DRG following traumatic nerve injury and during natural pain resolution. We provide a reference dataset comprising ∼7,500 tile-microscopy and confocal bioimages, along with a workflow demonstrating how objective, large-scale bioimage analysis can reveal sex-specific cellular differences in the DRG. To independently support the bioimage data, we provide mRNA transcriptome profiles from DRGs of both sexes. Our data show pronounced sex-specific molecular and cellular phenotypes which are most pronounced during natural, ongoing pain resolution.

## RESULTS

### Experimental strategy

We modeled pain resolution in adult male and female rats using the chronic constriction injury (CCI) model of the sciatic nerve (Figure 1). A priori, we defined the onset of pain resolution as the time point at which mechanical hypersensitivity, assessed by the von Frey test, was reduced by 50% relative to peak sensitivity (Figure 1A). This 50% threshold was selected because a similar magnitude of pain reduction is considered clinically meaningful by patients (31). Consistent with previous work from our group (32), peak mechanical hypersensitivity occurred within two weeks post-CCI, and the 50% recovery threshold was consistently reached by five weeks in both sexes, with no significant sex differences (Figure S1). For cellular phenotyping, single DRGs from lumbar segments L4 and L5 (16) were harvested and processed for systematic bioimage analysis, with a focus on sensory neurons, SGCs, and local macrophages. DRG mRNA transcriptomes were determined in both sexes at 1 and 5 weeks after injury from the ipsi- and contralateral side, in total 50 mRNA transcriptomes (Figure 1B). In addition, we also sequenced mRNA from male DRG 24h after injury (6 ipsilateral, 6 contralateral) for an initial injury phenotype. Statistical analyses assessed injury-, resolution-, and sex-specific differences (Figures 1C–E).

**Figure 1.**
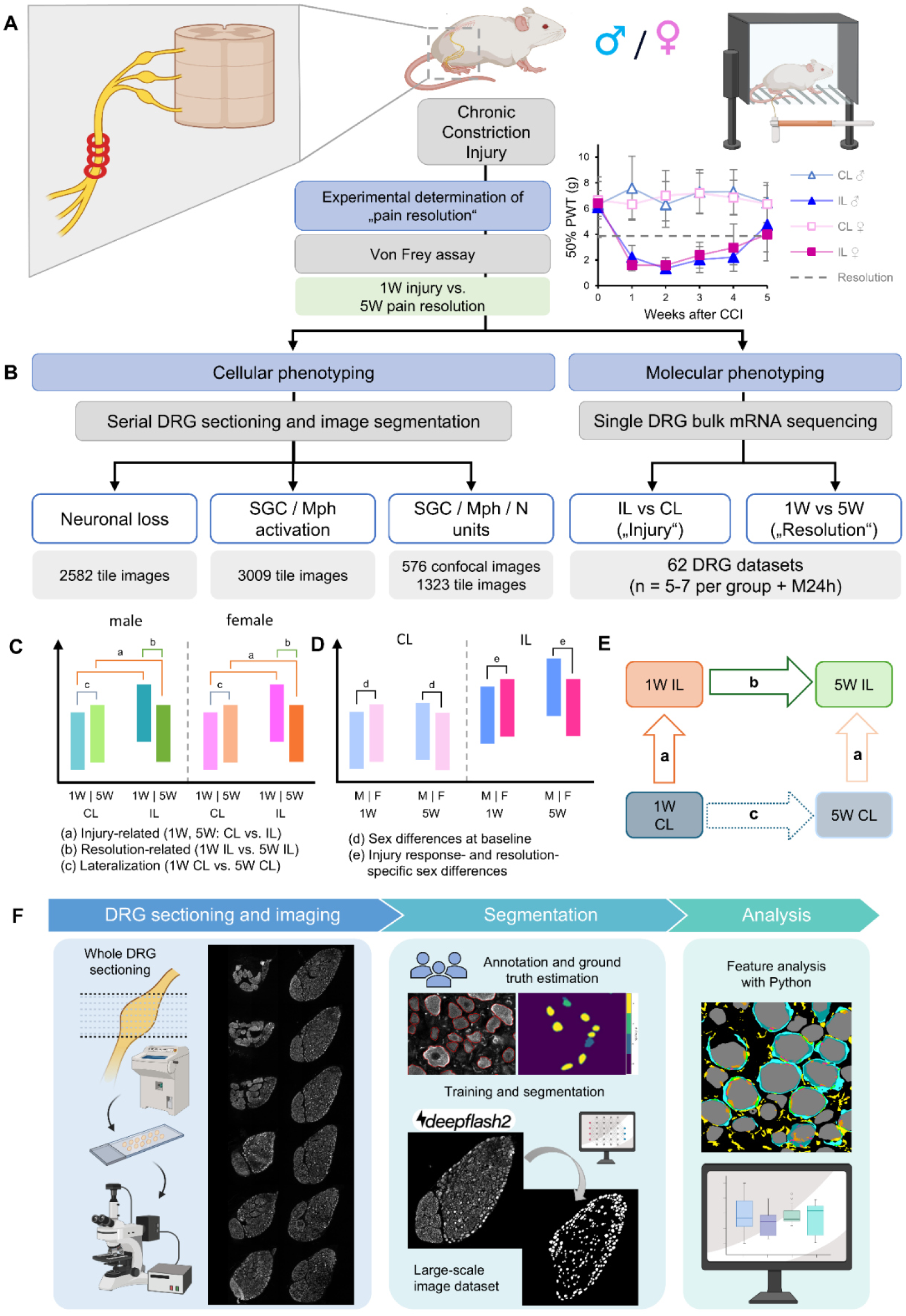
**Study design and bioimage analysis workflow.** (A) Ongoing pain resolution was modelled in male and female rats after CCI of the sciatic nerve. 50% recovery from mechanical hypersensitivity in the von Frey test was defined à priori to determine an objective time point for ongoing pain resolution (data in Figure S1). (B) Cellular phenotyping on bioimages was used to investigate neuron abundance, SGC and macrophage (Mph) phenotypes, and changes in the spatial organization of the three cells types close to the sensory neurons. The number of analyzed tile images and confocal images is indicated. Molecular phenotyping occurred based on deep bulk mRNA sequencing (RNA-Seq) data from single DRG. (C-D) Hypothesis testing for bioimage analysis. (C) Changes related to the injury (ipsilateral (IL) versus contralateral (CL)) at one week (1W) or five weeks (5W) after CCI, resolution of pain, and stage-dependent changes on the contralateral side (lateralization) were tested. (D) Changes related to sex were tested by comparing side-specific changes (CL / IL) between males and females, after injury (1W), and during ongoing pain resolution (5W). (E) Sample groups and comparisons included in the transcriptome analysis. (F) Workflow of whole DRG sectioning to image analysis with deepflash2. Serial sectioning, immunolabeling, and tile-microscopy was performed on entire DRG sections, across 10 microscope slides per DRG. Immunolabels were independently annotated by three experts to compute a consensus segmentation dataset. After ground truth estimation and training of DL model ensembles, neurons and SGCs were segmented. The raw images and segmentation masks were organized in a large-scale bioimage dataset. Bioimage feature analysis was finally computed with Python scripts (see also Figure S2 and Figure S3).

To ensure validity of the bioimage analysis, we used a deep learning (DL)-based image feature segmentation strategy, implemented via *deepflash2* (13, 16, 33, 34) (Figure 1F, Figure S2). For the neurons, we trained the DL model for image feature segmentation of the somata. Neurons and SGCs were segmented using DL model ensembles, achieving dice similarity scores ≥0.85 on both validation and test image sets (Tables S1, S2, S3). Tissue area and macrophages, identified with Iba1 (13), were segmented using advanced thresholding (Figure S3).

This way, we analyzed a total of 6,914 tile microscopy images, each consisting of 4-10 stitched sub-images covering entire DRG cross sections. The 6-16 slices per DRG contained on average 1,500-2,500 neurons in a tissue area of 7-13 mm^2^ per DRG for each staining (Figure S5). High resolution spatial analysis of the neuron-SGC and macrophage units was done on 576 confocal images. The bioimage dataset is publicly available via the BioImage Archive (https://www.ebi.ac.uk/biostudies/bioimages/studies/S-BIAD1944; DOI: 10.6019/S-BIAD1944). The images and respective segmentations can be viewed online (https://drg-pain-resolution.streamlit.app/).

### Tissue size and neuron numbers in rat DRG are sex-specific

First, we looked at basic system parameters like DRG tissue area and sensory neuron numbers in the 2D bioimage data. We determined the neuronal DRG area based on NF labeling and used a DL model to segment the somata of NF^+^ sensory neurons (Figure 2A). The tissue area for each slice was variable, as expected (Figure S4A), but the mean tissue area per slice analysis indicated that the DRG of males is bigger at 5 weeks than at the 1-week timepoint (Figure S4B). To exclude the possibility of variability of the tissue area due to the cutting angle, we calculated the sum of the total area captured per DRG. In males but not females, the total tissue area increased significantly between 1 and 5 weeks after CCI on the ipsilateral and contralateral sides (Figure 2B). Accordingly, the neuron density time-dependently decreased on the ipsilateral and contralateral side in males, but not females (Figure 2C). The neuron number per slice remained constant in both sexes over time and after injury (Figure 2D). To address a potential selective loss of NF^+^ neurons, we quantified neurons in 13 size groups (<200–2,600 µm²) (Figure 2E). Across all groups – sexes and time points – neuron numbers remained comparable, with only minor changes between ipsilateral and contralateral in the largest neurons. There was a linear correlation between the number of NF^+^ neurons per slice and the corresponding tissue area, but the variability of this measure of neuron density was remarkable high (Figure 2G).

**Figure 2.**
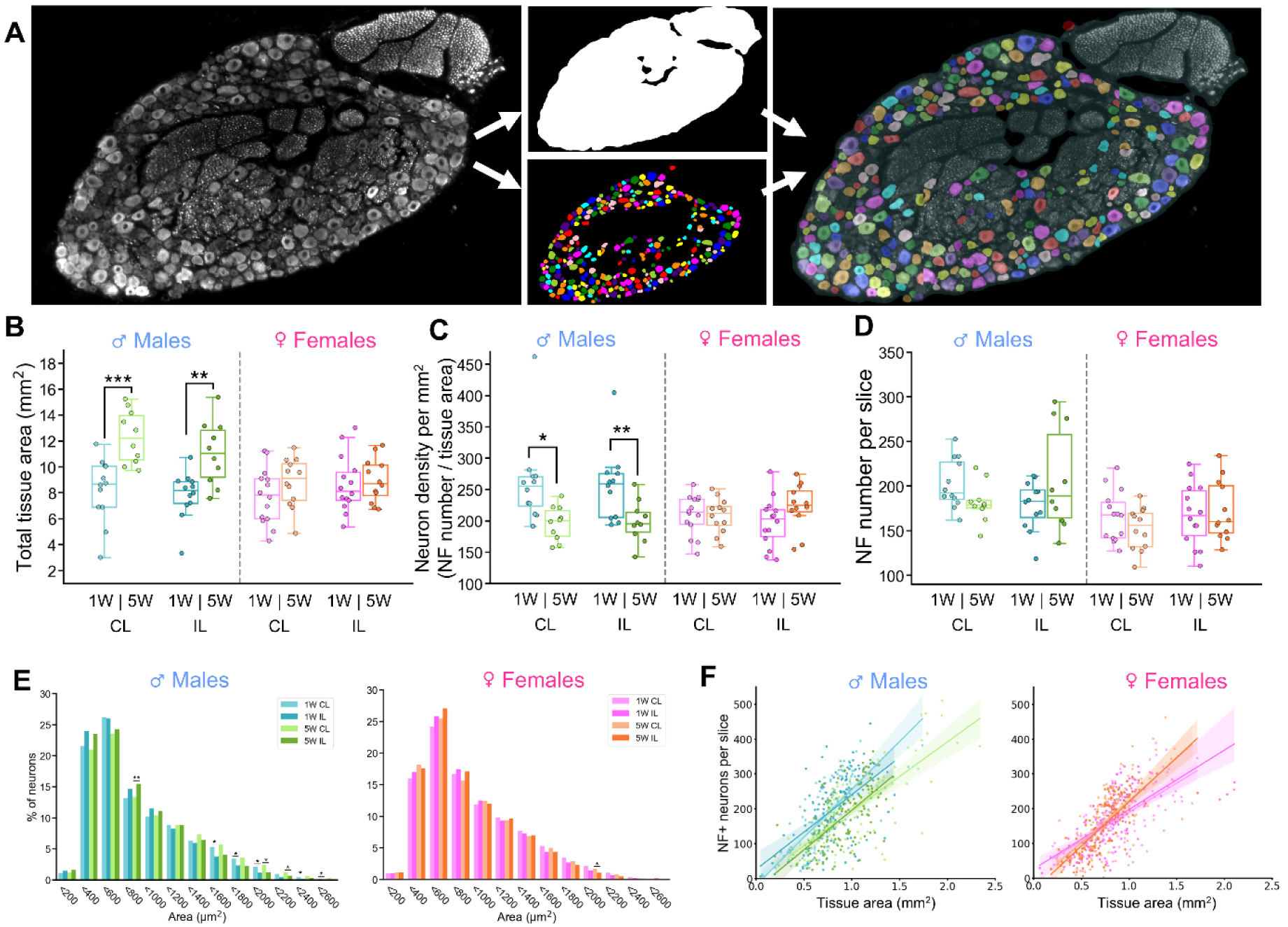
**DRG tissue growth and abundance of NF^+^ neurons after CCI.** Data of tissue size and neuron number analysis are shown for male (blue/green) or female (pink/orange) rats, 1W/5W after CCI. (A) Representative tile image of an anti-NF immunofluorescence staining (left) and the corresponding segmentation masks (middle). The NF staining was segmented for neuronal tissue area (white) and single sensory neuron cell bodies (color coded). Right: merge. (B) Total tissue area (mm^2^) as sum of the area of all 2D slices per DRG. (C) Mean neuron density (neurons per mm^2^ neuronal tissue area). (D) Mean number of neurons per slice in each DRG. (E) Size distribution of neurons in male (blue/green) and female (pink/orange) rats. Neurons were divided into bins of 200 µm^2^ cell area. The percentage of neurons in each bin compared to the total neuron count is shown. (F) Graphs showing the correlation between the tissue area and the number of neurons with single slice resolution (single data points) in male (blue/green) and female (pink/orange) rats. In (B-D), each data point represents one DRG (two DRG per side per animal), number of animals: female 1W, n = 7; male 1W, n = 6; female 5W, n = 6; male 5W, n = 5. P-values: *p ≤0.05; **p ≤0.01; ***p ≤0.001. Significance was tested for 1W CL vs. 1W IL, 5W CL vs. 5W IL (“Injury-related changes”), 1W IL vs 5W IL (“Resolution-related changes”), and 1W CL vs. 5W CL (“Lateralization of effects”), separate for both males and females (Bonferroni corrected per hypothesis). In E, statistical significance was tested for Injury-related changes only. In F, number of slices analyzed: female, 1W IL, n = 142; 1W CL, n = 134; 5W IL, n = 140; 5W CL, n = 144. For the males, 1W IL, n = 132; 1W CL, n = 122; 5W IL, n = 108; 5W CL, n = 132.

A recent publication states that non-peptidergic, small diameter neurons of the mouse, which can shape mechanical inflammatory hypersensitivity (35), are particularly susceptible to loss after traumatic sciatic nerve injury (14). To assess whether CCI leads to loss of this population, we analyzed DRG sections stained for NF, IB4, and ATF3 (Figure 3.A). Segmentation confirmed IB4⁺ labeling in small-diameter sensory neurons (Figure 3B). In both sexes, the number of IB4^+^ neuron labels per slice (Figure 3C) was reduced 1 week after CCI and in females also at 5 weeks after CCI and when normalized to the tissue area (Figure S5A). When we normalized the IB4⁺ labels to all NF⁺ neurons, the ratio of IB4^+^ cells was reduced by the injury under all conditions and in both sexes (Figure 3D). Furthermore, counting the total number of IB4^+^ cells confirmed a reduction of the label by the injury (Figure S5B). Especially IB4^+^ small diameter neurons with 200-600 µm in diameter were less abundant at the ipsilateral side at both 1 week and 5 weeks after CCI (Figure 3E). A slice-by-slice analysis showed a linear but highly variable number of IB4⁺ cells per individual slice (Figure 3F). However, since neither the NF^+^ sensory neuron segmentation, nor the mRNA-Seq data (see below), confirmed a gross loss of sensory neurons, we conclude that the nerve injury reduced IB4 immunoreactivity in the DRG of both sexes.

**Figure 3.**
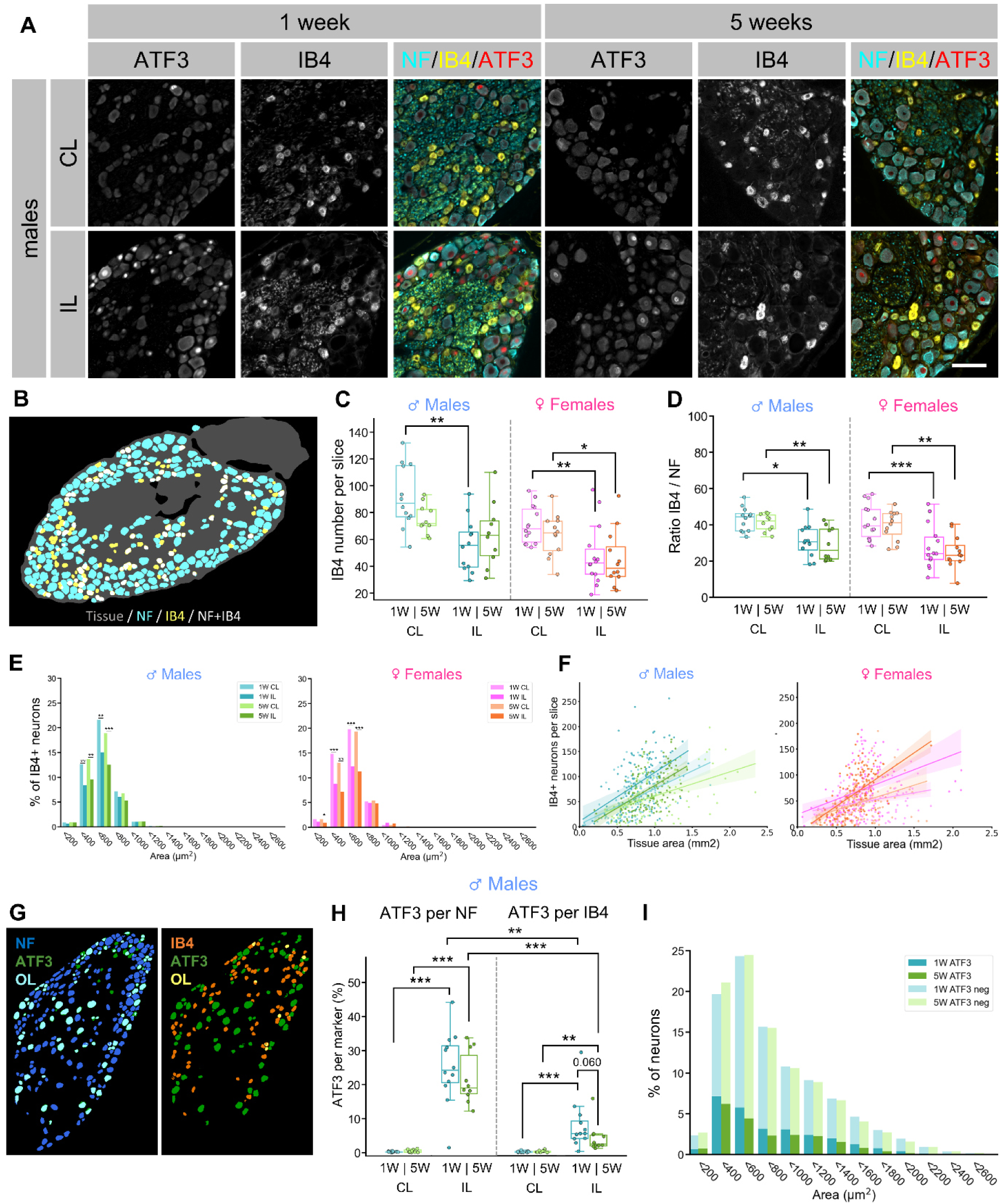
**Chronic constriction injury does not induce neuronal loss, but reduces IB4-marker abundance in male and female rats.** (A) Immunofluorescent tile images of rat DRG (L5), representing neurofilament heavy chain-positive (NF^+^) neurons, Isolectin B4-positive (IB4^+^) non-peptidergic neurons, and injury marker ATF3^+^ neurons 1 or 5 weeks (1W, 5W) after CCI. Scale bar: 100 µm. (B) Exemplary depiction of an overlay of the computed segmentations for the tissue area (gray, NF-based), NF^+^ neurons (cyan), IB4^+^ neurons (yellow) and their overlap (white). (C, D) Box plot showing the number of IB4^+^ neurons per slice (C), and their ratio per NF+ neurons (D) in male (blue/green) or female (pink/orange) rats, 1W/5W after CCI. (E) Size distribution of IB4^+^ neurons and in male (blue/green) and female (pink/orange) rats. Neurons were divided into bins of 200 µm^2^ cell area. The percentage of IB4^+^ neurons in each bin compared to the total neuron count (NF^+^) is shown. (F) Graphs showing the correlation between the tissue area and the number of IB4^+^ neurons with single slice resolution (single data points) in male (blue/green) and female (pink/orange) rats. (G) Exemplary depiction of an overlay of the segmentations of ATF3 (green) with NF (blue, left panel) and their overlap (OL, cyan), or IB4 (orange) and their overlap (OL, yellow), respectively. (H) Boxplots showing the percentage of NF^+^ neurons or IB4^+^ neurons with ATF3^+^ neurons in male rats, one (blue) and five weeks (green) after CCI. (I) Size distribution of ATF3^+^ and ATF3^-^ neurons one (blue) and five (green) weeks after CCI, on the ipsilateral side. Neurons were divided into bins of 200 µm^2^ cell area. The percentage of ATF3^+^ or ATF3^-^neurons in each bin compared to the total neuron count (ATF3 positive and negative) is shown. In (B-D), each data point represents one DRG (two DRG per side per animal), number of animals: female 1W, n = 7; male 1W, n = 6; female 5W, n = 6; male 5W, n = 5. P-values: *p ≤0.05; **p ≤0.01; ***p ≤0.001. Significance was tested for 1W CL vs. 1W IL, 5W CL vs. 5W IL (“Injury-related changes”), 1W IL vs 5W IL (“Resolution-related changes”), and 1W CL vs. 5W CL (“Lateralization of effects”), separate for both males and females (Bonferroni corrected per hypothesis). In E, statistical significance was tested for Injury-related changes only. In F, number of slices analyzed: female, 1W IL, n = 142; 1W CL, n = 134; 5W IL, n = 140; 5W CL, n = 144. For the males, 1W IL, n = 132; 1W CL, n = 122; 5W IL, n = 108; 5W CL, n = 132. CL, contralateral; IL, ipsilateral; W = weeks; OL = overlap

We also asked how injury responses, marked by ATF3, were distributed in neuronal subpopulations. For this, we computed DL models to detect sensory neurons with ATF3 in the nucleus of the sensory neurons (Figure 3G), or both ATF3^+^ and ATF^-^ neurons. Ipsilaterally, 20–30% of NF⁺ neurons expressed ATF3 (Figure 3H, left panel), significantly more compared to ∼8% of IB4⁺ neurons (Figure 3H, right panel). Although IB4⁺ ATF3 activation tended to be lower at 5 weeks than at 1 week, this change was not statistically significant. ATF3⁺ neurons spanned all size categories (Figure 3J). For experimental reasons (discussed later), ATF3 segmentation was only reliable in males and was therefore not plotted for females.

In summary, neuron number estimates in both sexes are challenged by sex-specific parameters of tissue size and neuron numbers, age-related growth of the DRG, as well as high variability in neuron distribution. Transient nerve injury did not induce overt loss of sensory neurons within 5 weeks post-injury. Instead, rat males showed age-related DRG tissue expansion that reduced the neuronal density due to more non-neuronal tissue.

### SGCs and macrophages show sex-specific phenotypes during pain resolution

Peripheral nerve injury induces plastic changes in SGCs and local macrophages at the sensory neuron border (16, 36). To test for non-neuronal cell phenotypes, we labeled for GFAP, which is specifically upregulated in SGCs after injury (16), and Iba1 (allograft inflammatory factor; *Aif1*) to label for macrophages close to the neuron border (13) (Figure 4A). Since macrophages cluster near neurons after injury (37, 38), we asked how this affects the spatial distribution of the SGCs. To define the space close to the neurons, we computed a neuron-near area based on the somatic segmentation of the NF label (Figure 4B).

**Figure 4.**
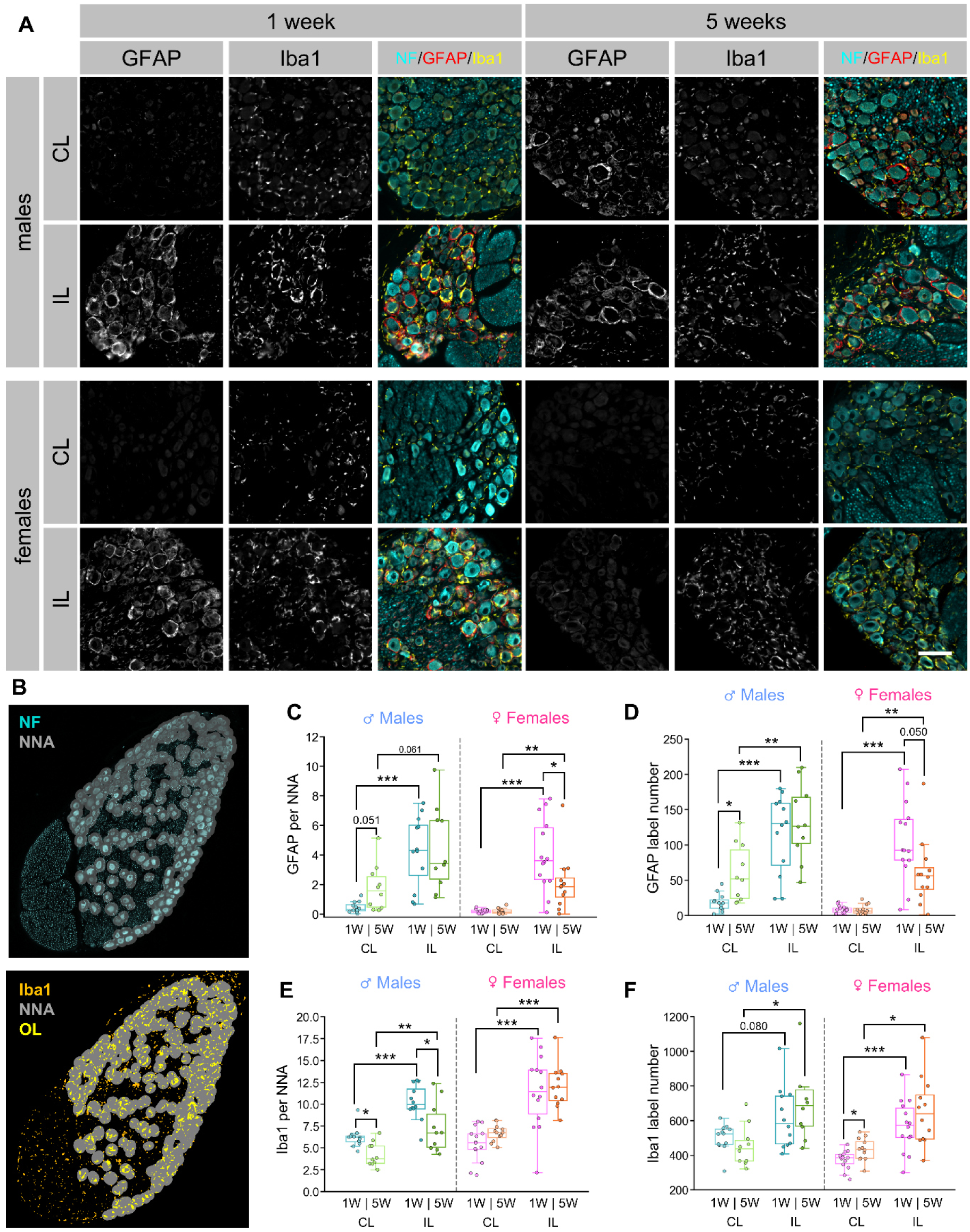
**Activation of satellite glia persists in males only, whereas macrophage activation persists in females only.** (A) Immunofluorescent tile images of rat DRG (L5), representing neurofilament heavy chain-positive (NF^+^) neurons, GFAP^+^ activated SGC, and Iba1^+^ macrophages one or five weeks after CCI. Scale bar: 100 µm. (B) Example of computed neuron-near area (NNA, grey) as dilation of the NF segmentation, overlaid over the corresponding NF staining (cyan) (upper panel) or Iba1 staining (yellow) (lower panel). (C-F) Box plots showing the mean GFAP^+^ area per NNA per slice (C), mean number of distinct GFAP labels per slice (D), mean Iba1^+^ area per NNA per slice (E), and mean number of distinct Iba1 labels per slice (F) for male (blue/green) or female (pink/orange) rats, 1W/5W after CCI. Each data point represents one DRG (two DRG per side per animal), number of animals: female 1W, n = 7; male 1W, n = 6; female 5W, n = 6; male 5W, n = 5. P-values: *p ≤0.05; **p ≤0.01; ***p ≤0.001. Significance was tested for 1W CL vs. 1W IL, 5W CL vs. 5W IL (“Injury-related changes”), 1W IL vs 5W IL (“Resolution-related changes”), and 1W CL vs. 5W CL (“Lateralization of effects”), separate for both males and females (Bonferroni corrected per hypothesis). CL, contralateral; IL, ipsilateral; W = weeks; NRA = neuron-rich area; OL = overlap

In both sexes, GFAP immunoreactivity in SGCs was significantly increased ipsilaterally 1 week after CCI, while contralateral DRGs were largely GFAP-negative (Figure 4C, D). In females, GFAP levels significantly declined between 1 week and 5 weeks post-CCI in the neuron-near area. In contrast, GFAP immunoreactivity persisted throughout the 5-week period in the males (Figure 4C, D). In males, there was also a trend toward increased contralateral GFAP at 5 weeks (Figure 4C). These patterns were confirmed by counting GFAP^+^ “rings” (label number) surrounding sensory neurons (Figure 4D). As expected, more Iba1^+^ macrophage labels were found ipsilaterally at 1 week and 5 weeks post-CCI in the neuron-near area, in both sexes (Figure 4E). In females, elevated Iba1 levels persisted through the pain resolution state, whereas in males Iba1^+^ labels were significantly reduced in the neuron-near area, both, on the ipsilateral and contralateral side. Note that the pure number of Iba1 labels did not change on the injury side over time (Figure 4F).

To better understand the cellular changes in the neuron-near area, we quantified the distribution of neurons (NF^+^), SGCs (FABP7^+^), and macrophages (Iba1^+^) at higher resolution with confocal imaging of DRG sections (Figure 5A). We used FABP7 for this experiment, as the protein is highly expressed in SGCs even in the absence of injury (39). Moreover, FABP7 labels clearly outline the SGC soma, allowing reliable separation of SGCs and local macrophages by cell segmentation (13). In females and males, on the ipsilateral side, we saw lower FABP7 abundance at 5 weeks after CCI (Figure S6A) and a higher Iba1 label abundance after injury in both sexes, as expected (Figure S6B). The overlap between both labels, FABP7 and Iba1, increased after the injury and decreased again during resolution (Figure S6C), indicating spatial dynamics in both cell types.

**Figure 5.**
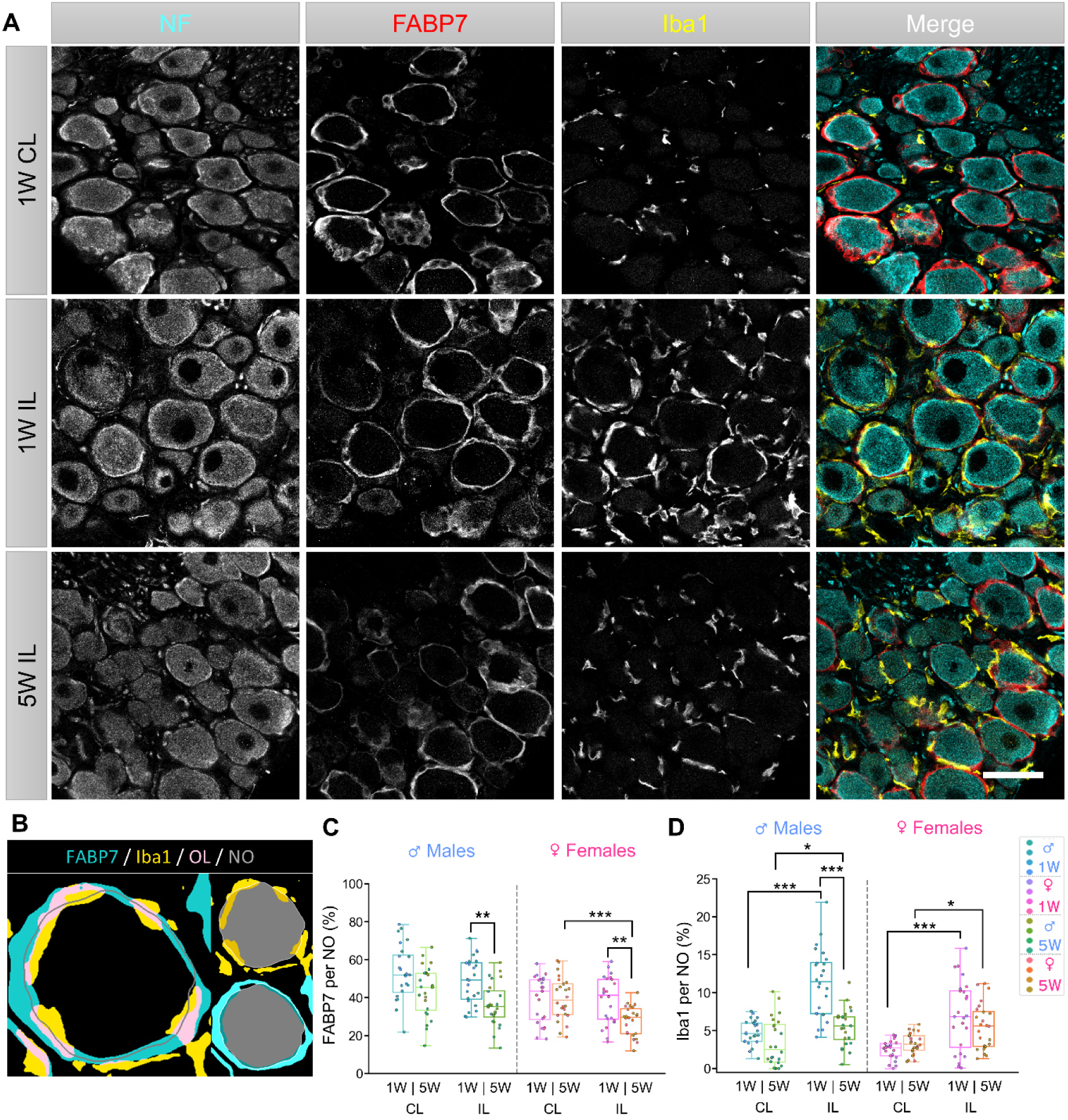
**Neuronal injury-dependent increase of macrophage-neuron contact but long-term reduction of SGC-neuron contact.** (A) Confocal images of rat DRG (L5) stained with neurofilament heavy chain (NF, cyan), SGC-marker FABP7 (red) and macrophage marker Iba1 (yellow). Scale bar 50 µm. (B) Example of segmentations of one neuron and its border in contact with either FAPB7^+^ SGCs or Iba1^+^ macrophages (NO = neuron segmentation outline; OL = overlap). (C,D) Box plots show the quantification of FABP7^+^ (C) and Iba1^+^ (D) staining per neuron segmentation outline. (n = 24 images per condition, with 6 images of 4 rats). Significance is indicated as follows: *p ≤0.05; **p ≤0.01; ***p ≤0.001. Significance was tested for 1W CL vs. 1W IL, 5W CL vs. 5W IL (“Injury-related changes”), 1W IL vs 5W IL (“Resolution-related changes”), and 1W CL vs. 5W CL (“Lateralization of effects”) and Bonferroni corrected per hypothesis. CL, contralateral; IL, ipsilateral; W = weeks;

Next, a DL model was trained to segment the neuron border for precise calculation of SGC and macrophage contacts with the neurons (Figure 5B, NO = neuron segmentation outline). In both sexes, FABP7 contact with the neuron border was significantly reduced ipsilaterally over pain resolution, from 1 week to 5 weeks. In females, we still found an injury-related effect at 5 weeks (CL compared to IL; Figure 5C). In contrast, for macrophages, the contact with the neuron border was significantly increased ipsilaterally after injury (Figure 5D). Of note, this was significantly reduced only in males during the pain resolution phase (Figure 5D).

Recently, using glutamine synthetase labeling (GS; = Glul, glutamate ammonia ligase), we reported phenotype plasticity of SGCs after spared nerve injury (16). Furthermore, reports confirmed high abundance of GS in different SGC clusters in single cell categorization experiments (40, 41). Therefore, we asked whether the SGC and nearby macrophages show spatial dynamics close to injured neurons, in particular. We labeled GS to look for SGC plasticity, Iba1 for local macrophages and ATF3 to label injured neurons (Figure S7). Basal somatic levels of ATF3 also let us delineate uninjured neurons. Therefore, we trained two different DL models on the ATF3 labels, one to detect the neurons (ATF3all) and one to detect neurons with ATF3 activation (ATF3-positive). From the neuron segmentations, we computed the neuron outline for each neuron and its overlap with either GS or Iba1 labels (Figure S7B). Compared between 1 week and 5 weeks post-CCI, the abundance of GS-positive cells close to the neuron border was reduced, ipsi- as well as contralateral (Figure S7C, D). Then, we focused on the ipsilateral side and computed GS abundance close to ATF^-^ (somatic ATF3, but no nuclear ATF3) neurons in comparison to ATF3^+^ neurons. This analysis showed that GS abundance close to neurons is independent of whether individual sensory neurons show ATF3 activation (Figure S7E). In the same dataset, we could also confirm the above-mentioned macrophage dynamics in male rats after injury at 1 week (Figure S7F, G) and additionally saw a higher overlap of macrophages with the ATF3^+^ versus ATF3^-^ neuron outlines at 5 weeks after injury (Figure S7H).

The data reveal that CCI induces sex-specific and time-dependent changes in both SGCs and macrophages. SGC phenotyping is challenging due to sex- and time-dependent effects observed for the markers FABP7, GS, as well as GFAP. After injury, SGCs are displaced from the neuron border and macrophages become more abundant in the space between neurons and the SGCs. During pain resolution, macrophages withdraw from this space, an effect more pronounced in males than in females. Nevertheless, macrophage activation after injury remains present during the pain resolution phase.

### Comparison between female and male cellular phenotypes

While we first tested differences within each sex, we also calculated the statistical differences between females and males (Figure S8). In short, the data indicate that males have a larger DRG tissue size at 5 weeks (Figure S8A, B). Females show slightly lower numbers of sensory neurons in the NF and IB4 stainings, but a similar ratio of IB4 to NF and the tissue area as males (Figure S8C-H). For SGCs, females display a lower abundance of GFAP activation, particularly during resolution, and a generally lower coverage of neurons by FABP7^+^ SGCs (Figure S8I-M). Overall, there is a higher abundance of macrophages at 1 week post CCI in males (Figure S8M-Q). In both sexes, the injury increases the abundance of macrophages directly at the sensory neurons, an effect which is significantly more resolved in males than females.

### Common transcriptome changes after CCI in both sexes

To obtain further experimental evidence of sex-specific changes during pain resolution, we performed mRNA sequencing of single DRG from female and male DRG (∼2 – 3 × 10^7^ reads per DRG, transcript per million (TPM) normalization: data table S1; DESeq2-normalization, data table S2).

First, we performed cell-type marker analysis based on the harmonized DRG atlas (40) (Figure S9). We observed abundant expression of non-neuronal cell markers for SGCs, Schwann cells, macrophages, neutrophils, pericytes, fibroblasts, and endothelial cells (Figure S11A). Among these, macrophage markers (*Cd74, Csfr1, Cx3cr1, Itgam (CD11b)*) were significantly upregulated after injury, consistent with the macrophage activation phenotype observed above. SGC activation marker *GFAP* was also upregulated, in agreement with the immunolabeling data. During the resolution phase, females showed a modest downregulation of Schwann cell markers (*Cadm2, Mbp, Mpz*), whereas males exhibited slight decreases in *Fabp7* and *Gfap* in SGCs (Figure S9A). *GS* (*Glul1*) was virtually unchanged in the RNA data.

Immunohistochemical cellular phenotyping had revealed no gross neuronal loss, though anti-IB4 immunoreactivity was significantly reduced. In the transcriptome data, no evidence of loss was observed for subtype-specific neuronal markers. In particular, non-peptidergic neurons (NP, typically IB4^+^) retained transcripts encoding *Mrgprd, Hrh1, Scn11a (Na_V_1.9)*, and others. By contrast, injury-associated genes such as *Atf3* and *Gal* were strongly upregulated post-injury (Figure S9). Pan-neuronal markers remained comparable across all conditions. These findings support the conclusion drawn from cellular phenotyping that CCI in rats does not induce neuronal loss within five weeks post-injury.

Across conditions, more genes were upregulated than downregulated on the injury side (Figure 6A). Volcano plots highlighted *Atf3* and the neuropeptide *Npy* as the most strongly regulated genes in males, whereas *Atf3* and *Stac2*—a regulator of calcium channel activity—were most prominent in females. At the pain-resolution stage (5 weeks), the number of DEGs was reduced, yet many overlapped with those observed at 1 week. The top 25 DEGs were visualized in heatmaps (Figure 6B), revealing consistent upregulation of regeneration-associated transcription factors (*Atf3*, *Sprr1a*), neuropeptides (*Npy*, *Gal*, *Vip*), cytokines (*Il6*, *Il24*), adhesion-related genes (*Cldn4*, *Smagp*), and species-specific injury response genes (*Csrp3*, *Stac2*, *Hamp*). Notably, *Sprr1a*, *Npy*, *Gal*, and *Csf1* — all strongly induced after CCI — are transcriptional targets of *Atf3* (42).

**Figure 6.**
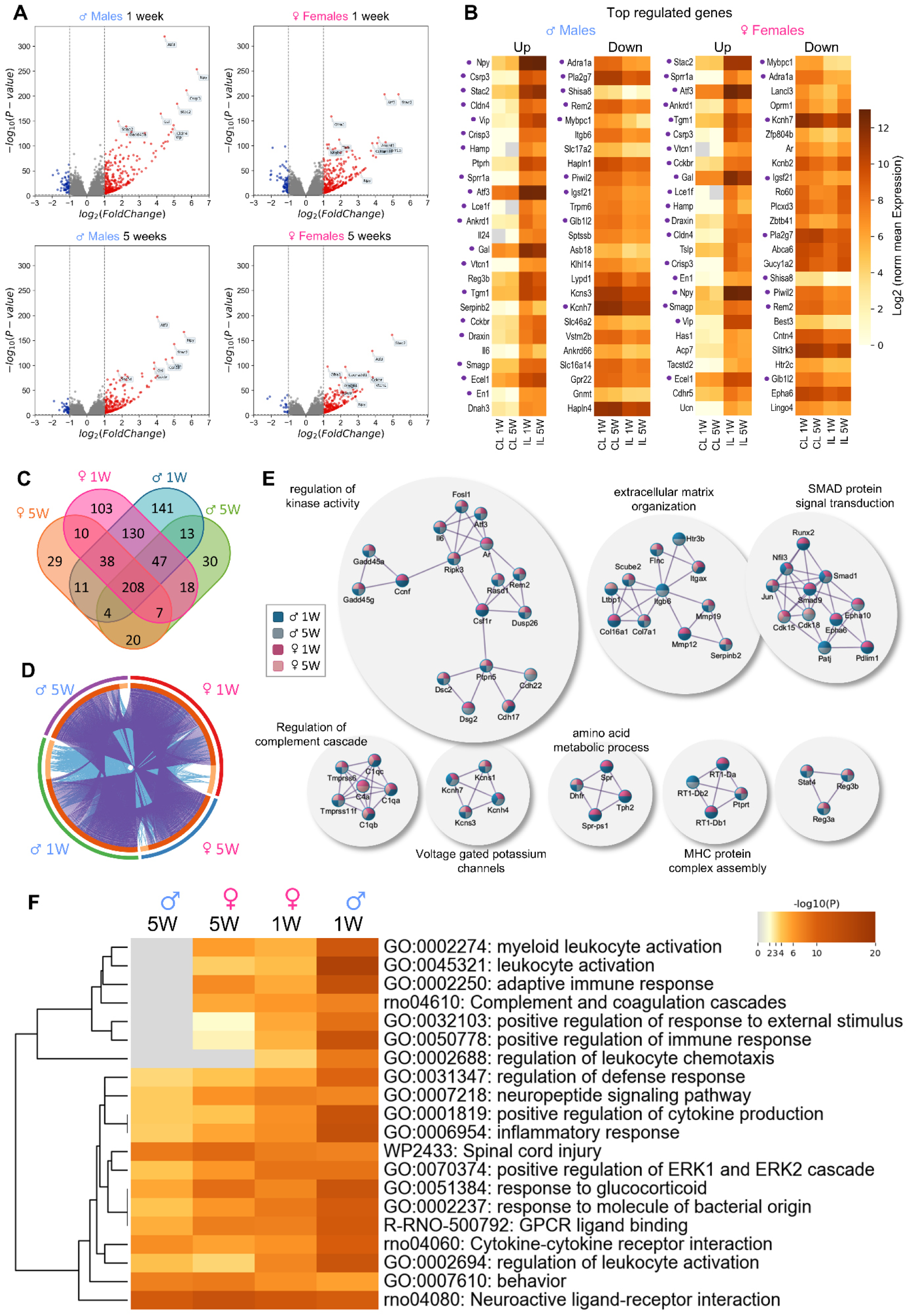
**Transcriptome analysis after chronic constriction injury shows a strong immune phenotype and regulation of signaling mediators.** All figures in this list are based on comparing transcriptome data from the ipsi- (IL) with the contralateral (CL) side, at 1 or 5 weeks (1W, 5W) after CCI in males or females. (A) Volcano plots highlighting the most significantly upregulated (red) and downregulated (blue) genes. Genes with an absolute log2fold change ≥ 1 and an adjusted p-value <0.01 were considered significantly differentially expressed. (B) Heatmaps illustrating the top upregulated (left panel) and downregulated (right panel) genes in males and females 1W after CCI. Genes that are in the top 25 regulated genes both in males and females are marked with a purple dot. (C) Venn diagram of shared and unique significantly differentially regulated genes for each condition. (D) Circos plot (https://circos.ca/) of overlaps between gene lists of significant differential regulation. Purple lines link identical genes between conditions and blue lines link genes with shared ontology terms. Created with metascape.org. (E) Protein-protein interaction (PPI) enrichment analysis of male and female significantly differentially regulated genes, created with metascape.org and modified with cytoscape.org. PPIs are clustered with the Molecular Complex Detection (MCODE) algorithm and best scoring terms are indicated next to the clusters as functional descriptors of the components. Proteins that appear in male (blue) or female (pink) lists are color-coded as dark (1W) or light (5W), respectively. (F) Heatmap of enriched terms in pathway analysis coded by p-value, created with metascape.org. n = 7 (M, F 1W), n = 6 (F 5W), n = 5 (M 5W). CL, contralateral; Il, ipsilateral; W = weeks; M = male, F = female; DE = differentially expressed.

Downregulated genes were more heterogeneous between sexes; of the top 25 DEGs, only nine overlapped between males and females (*Adra1a*, *Pla2g7*, *Shisa8*, *Igsf21*, *Mybpc1*, *Rem2*, *Glb1l2*, *Piwil2*, *Kcnh7*). Several were associated with synapse and nervous system development (*Hpln1*, *Rem2*), while others were linked to neuronal excitability (potassium channel genes *Kcnh7*, *Kcns3*, as well as *Trpm6* and *Rem2*).

A substantial fraction of DEGs (208 of 809) were shared across all injury conditions (CL vs. IL, 1 and 5 weeks, in males and females, Figure 6C, D). These genes were encoding, among others, for protein-protein interaction networks involved in *complement cascade regulation*, *kinase activity*, *extracellular matrix organization*, and *SMAD signaling* (Figure 6E, data table S3). Gene ontology analysis indicated that male rats exhibited a stronger early upregulation of immune-related pathways at 1 week, that was absent or reduced by 5 weeks, compared to a more stable upregulation in females (Figure 6F, data table S4). This was in line with more prevalent macrophage activation observed in the immunolabeling data. Together with our cellular phenotyping data, these results support a model of sex-specific macrophage functions after injury and during pain resolution.

For the males, we also checked the transcriptome at 24h after CCI in order to test for early transcriptome changes after injury. This dataset confirmed high induction of *Atf3* (neuronal) and *Gfap* (SGC) due to the injury and revealed early upregulation of signaling mediators such as *Alkal2, Il6, Vgf*, and the macrophage attractant *Ccl2* (Figure S10A-C). Gene ontology analysis pointed to regulation of the MAPK cascade and strong activation of immune phenotypes (Figure S10D, data table S5).

### Common and sex-specific transcriptomic changes in the DRG during pain resolution

To identify biological processes associated with pain resolution, we compared ipsilateral DRGs at 1 week and 5 weeks after CCI in both sexes. Similar numbers of genes were up- and downregulated (Figure 7A), indicating that resolution involves both activation and suppression of transcriptional programs.

**Figure 7.**
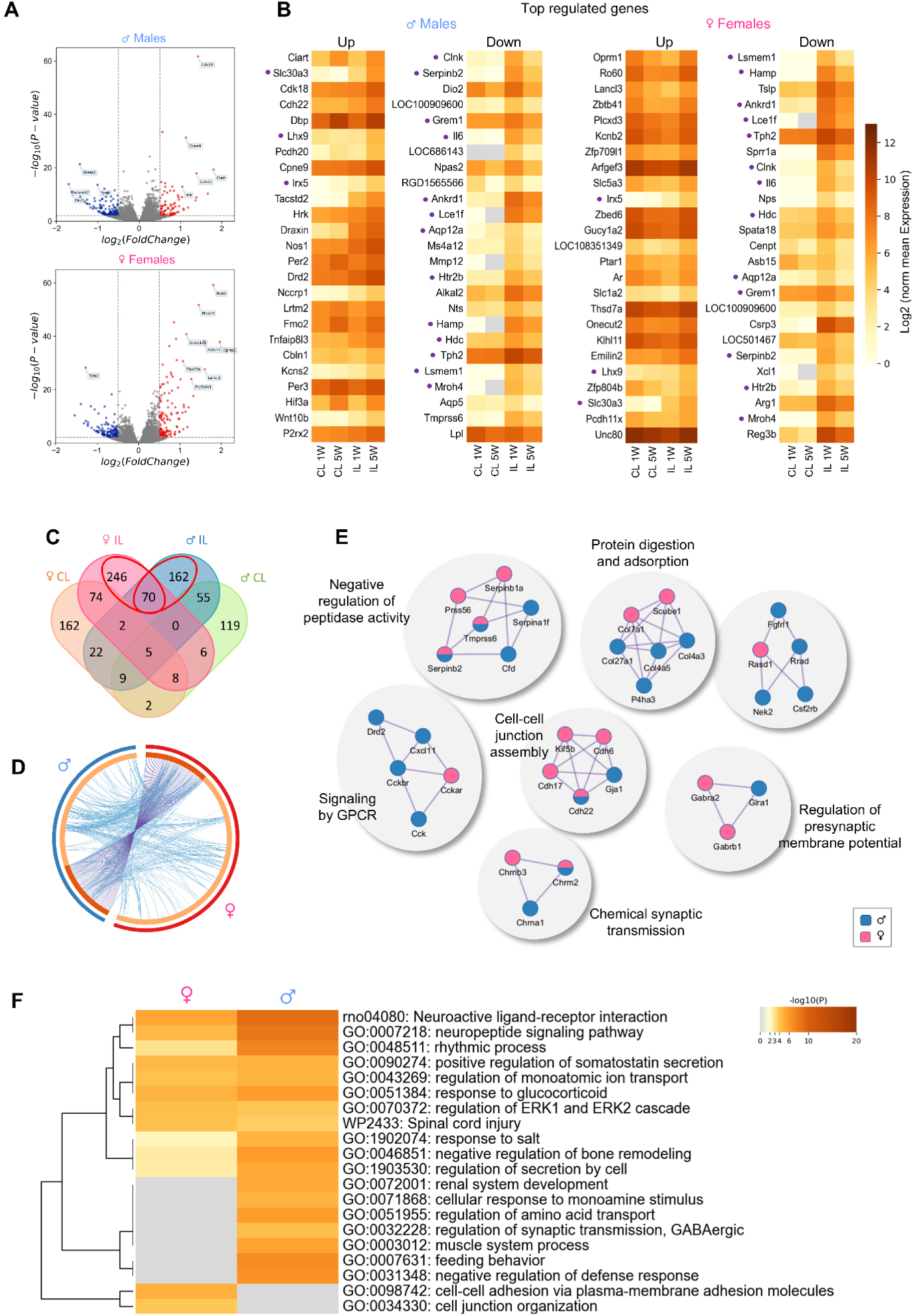
**Pain resolution is marked by sex-specific gene regulation patterns.** All figures in this list are based on comparing transcriptome data from the ipsilateral side between 1 and 5 weeks after CCI in males or females, respectively. (A) Volcano plots highlighting the most significantly upregulated (red) and downregulated (blue) genes. Genes with an absolute Log2fold change ≥ 0.5 and an adjusted p-value <0.01 were considered significantly differentially expressed. (B) Heatmaps illustrating the top upregulated (left panel) and downregulated (right panel) genes in males and females. Genes that are in the top 25 regulated genes both in males and females are marked with a purple dot. (C) Venn diagram of shared and unique genes for each condition, including a comparison of the contralateral sides. For further analysis, only genes that were specific to a change on the ipsilateral side were used (marked in red), to get strong, local effects on the ipsilateral side. (D) Circos plot (https://circos.ca/) of overlaps between gene lists of significant differential regulation. Purple lines link identical genes between conditions and blue lines link genes with shared ontology terms. Created with metascape.org. (E) Protein-protein interaction (PPI) enrichment analysis created with metascape.org and modified with cytoscape.org. PPIs are clustered with the Molecular Complex Detection (MCODE) algorithm and best scoring terms are indicated next to the clusters as functional descriptors of the components. Proteins that appear in male or female lists are color-coded as blue (M) or pink (F), respectively. (F) Heatmap of enriched terms in pathway analysis coded by p-value, created with metascape.org. n = 7 (M, F 1W), n = 6 (F 5W), n = 5 (M 5W). CL, contralateral; Il, ipsilateral; W = weeks; M = male, F = female; DE = differentially expressed.

When comparing the top 25 DEGs between sexes, only three genes—*Irx5*, *Slc30a3*, and *Lhx9*—overlapped (Figure 7B). *Irx5* encodes a transcription factor that regulates potassium channel promoter activity. *Slc30a3* (encoding zinc transporter 3), and *Lhx9* (encoding Lim homeobox 9) are genes of unknown significance (GUS) in the field of research. In females, the opioid receptor *Oprm1* and the androgen receptor *Ar* were among the most strongly upregulated genes during resolution. Thirteen genes were shared in the top 25 downregulated genes (*Hamp*, *Ankrd1*, *Lce1f*, *Tph2*, *Clnk*, *Il6*, *Hdc*, *Aqp12a*, *Grem1*, *Serpinb2*, *Htr2b*, *Lsmem1*, *Mroh4*). Notably, *Il6*, which was robustly upregulated after CCI, was significantly reduced at 5 weeks post-CCI in both sexes.

Overall, fewer genes were shared between males and females during resolution compared to the CCI condition (77 common genes; 334 unique to females; 248 unique to males; Figure 7C, D). Nevertheless, several protein-protein interaction networks were shared between sexes (Figure 7E, data table S6), suggesting convergence on similar biological processes through distinct gene sets. These shared processes included *negative regulation of peptidase activity*, *protein digestion and absorption*, *cell–cell junction assembly* (more pronounced in females), *regulation of presynaptic membrane potential*, and *GPCR signaling* (more pronounced in males). Gene ontology comparisons revealed both common pathways (e.g., *neuropeptide signaling*) and sex-specific processes, such as *regulation of amino acid transport* in males and *cell junction organization* in females (Figure 7F, data table S7).

Next, we looked specifically into the differences between males and females at the time point of ongoing pain resolution (5 weeks after CCI) (Figure S11). Volcano plots using the data from the non-injured side show sex-associated genes (Figure S11A). Genes exclusively found in males only were *Eif2s3y*, *Kdm5d*, *Uty* (all Y chromosome-linked), and *Ddx3* (Chr. 13-linked) (Figure S11B). To get a set of genes that are differentially expressed because of the injury, but not at baseline, we filtered out genes that appeared on the contralateral side and tested for an overlap between genes that were differentially expressed during the injury (Figure S11C, D). Interesting, highly regulated candidates included *Col7a1* on the male-associated side, relating to extracellular matrix reorganization, and the opioid receptor *Oprm1* in females. Genes that were more strongly expressed in females were associated with processes relating to *ion transmembrane transport*, *chemical synaptic transmission*, *regulation of cell-matrix adhesion*, and *positive regulation of cell development*, whereas terms more strongly expressed in males included *neutrophil degranulation* or *defense* / *immune functions* (Figure S11E, data table S8).

### Pain resolution-specific processes are distinct between sexes

Since pain resolution involves biological processes related to both the onset and recovery of pain, we integrated the gene expression data from the initial CCI injury response and the subsequent resolution phase for both sexes (Figure 8). For the injury-related genes, we restricted the gene set to the contra- to ipsilateral comparison at 1 week after CCI for the injury response (Figure 8 A, indicated with ‘a’). Resolution-related genes were identified by comparing 1 week and 5 weeks after CCI of the ipsilateral side (Figure 8A, indicated by ‘b’). Regulated genes observed between 1 week and 5 weeks on the contralateral side were defined as lateralization effects and were removed from the gene list (Figure 8A, indicated by ‘c’). By this, we found 560 genes that were injury-specific, and 449 genes were resolution-specific; the intersection between both groups (170 genes) represents genes that showed a reversal to the initial expression level before the injury (Figure 8B). Finally, we defined six groups for gene subsets that were either still up- or down-regulated at 5 weeks (“continuous expression”), returning to baseline expression levels (“reversal back up / down”), or were only regulated in the resolution phase (“resolution-specific” genes) (Figure 8C).

**Figure 8.**
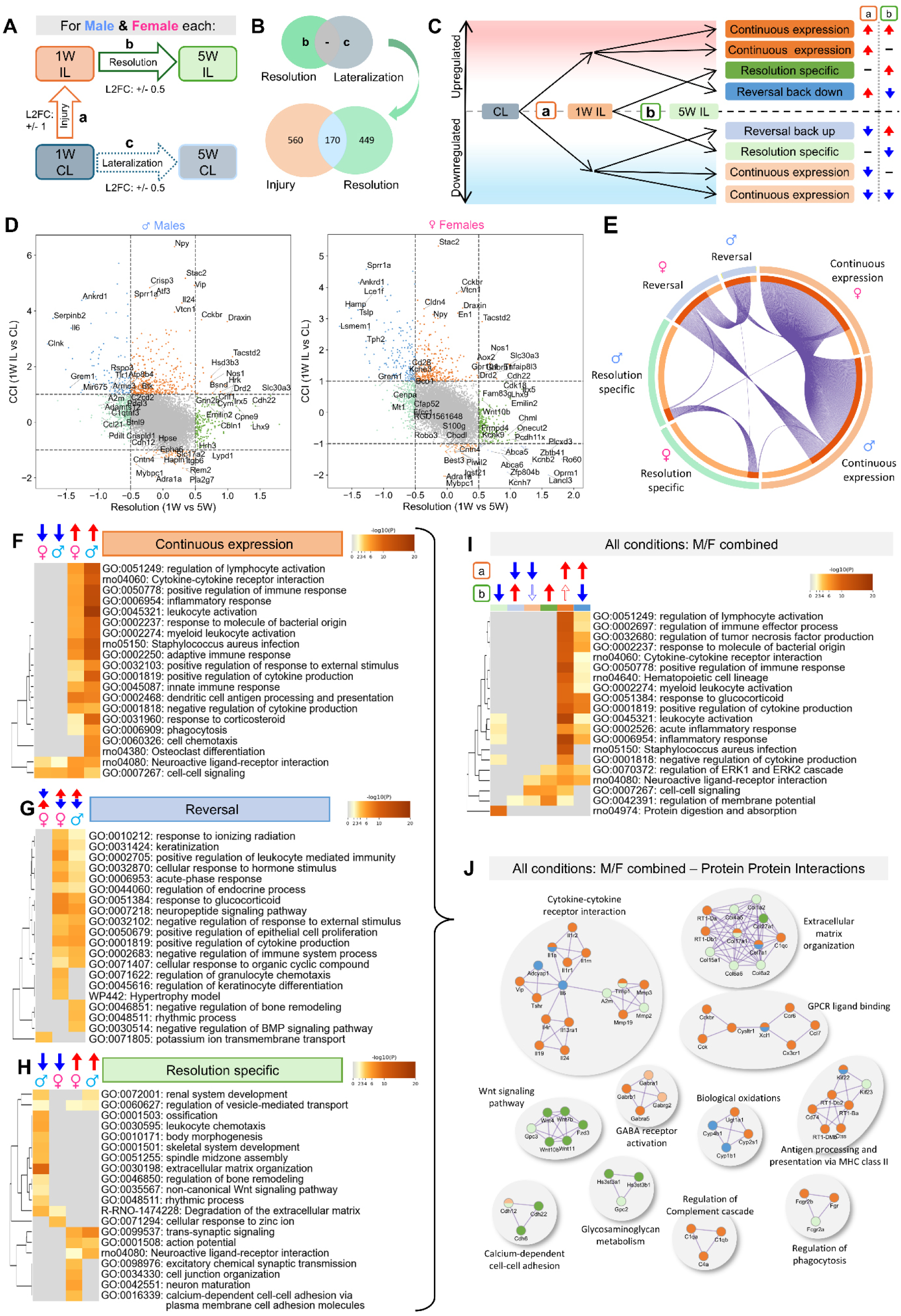
**Pain resolution is a sex specific process.** (A) Schematic depiction of gene expression comparisons used for this paper with their respective Log2fold change (L2FC) cutoff values. (B) Venn diagrams showing the steps of removing genes that lateralize (CL1W vs. CL5W) from the “resolution” gene pool before investigating overlap of the injury and resolution gene set. (C) Grouping of the gene comparisons according to their occurrence in gene set (a) injury and (b) resolution, taking up- (red) and downregulation (blue) into consideration, to extract groups of continuous expression (orange), resolution specific genes (green) and reversed expression (blue). Color scheme is maintained throughout the figure. (D) Scatterplot showing gene regulation groups based on the log2fold changes in the Injury (1WIL vs. 1WCL) and Resolution (5WIL vs. 1WIL) comparisons. (E) Circos plot (https://circos.ca/) of overlapping genes (purple lines) between male and female groups. Shown are differentially expressed genes (up and downregulated both) per group. Created with metascape.org. (F-H) Heatmap of enriched terms in pathway analysis with distinct groups for up- or downregulation and sex, for (H) continuous expression, (I) reversal (upper arrow injury regulation, lower arrow resolution), and (J) resolution specific genes. Coded by p-value, created with metascape.org. (I) Heatmap of enriched terms in pathway analysis of all groups (continuous, resolution, reversal), with distinction of up- and downregulation, M/F combined. Coded by p-value, created with metascape.org. (J) Protein-Protein Interaction (PPI) network of genes from all groups (continuous, resolution, reversal), with distinction of up- and downregulation, M/F combined. PPIs are clustered with the Molecular Complex Detection (MCODE) algorithm and best scoring terms are indicated next to the clusters as functional descriptors of the components. Created with metascape.org and modified with cytoscape.org.

Many genes, as *Il6* and *Ankrd1*, that were initially strongly regulated by the injury stimulus, were reversed in male and females (Figure 8D, upper left panel, blue). For males, there were almost no genes downregulated after injury and then regulated back up during pain resolution (“reversal back up” genes, Fig 8D, lower right panel, purple). For females, this category contained nine genes, including the opioid receptor *Oprm1* and the potassium ion channel proteins *Kcnh7* and *Kcnb2* (Figure 8D).

Continuously expressed genes included, *Npy* and *Stac2* in both sexes (Figure 8D, upper middle panel, dark orange). Also found in both sexes was the persisting downregulation of the alpha-1A adrenergic receptor (*Adra1a*) and the cell adhesion molecule contactin-4 (*Cntn4*) (Figure 8D, lower middle panel, orange). Resolution-specific genes included *Lhx9*, a Lim-HD protein involved in axonal guidance (43), or the extracellular glycoprotein *Emilin-2*, which is associated with extracellular matrix organization (Figure 8D, in green). Most resolution-specific genes were also sex-specific, as shown when comparing overlapping genes of each condition between male and female (Figure 8E). In contrast, genes in the reversal and continuous expression groups were mostly shared between both sexes.

Gene ontology analysis revealed that genes associated with *immune* / *inflammatory response* and *cytokine production* were still dominating the transcriptome profile (“continuous expression”). Genes associated with *neuroactive ligand-receptor interaction* and *cell-cell signaling* continued to be regulated in both directions (Figure 8F, data table S9). “Reversal“-associated genes were mostly reversed “back down” after initial upregulation in both sexes and involved genes associated with *neuropeptide signaling pathway*, *response to glucocorticoid*, and immune system regulation (Figure 8G, data table S10). In females, genes associate with *potassium ion membrane transport* were reversed back up, as described before. In the “resolution-specific” gene list, males showed a downregulation of genes associated with *extracellular matrix organization*, *non-canonical Wnt-signaling pathway* and *ossification* (Figure 8H, data table S11). In contrast, females showed resolution-specific upregulation of *neuron maturation* and *cell junction organization* genes. In both sexes, we saw a resolution-specific upregulation of genes involved in *trans-synaptic signaling* and *neuroactive ligand-receptor interaction* (also seen in “continuous expression”). Over all conditions, the continuous expression of genes associated with various immune system processes was the most prominent reaction to the CCI (Figure 8I, data table S12). Some of the immune system processes were reversed, and, thus, the “continuous expression” and “reversal” groups shared protein-protein interaction terms such as *cytokine-cytokine receptor interaction* (Figure 8J, data table S13). Resolution-specific processes were more distinct and sex-specific, including, for example, *calcium-depended cell-cell adhesion* in females and *Wnt signaling pathway* in males.

## Discussion

Sex is a key biological variable in neuropathic pain, yet the molecular and cellular processes that underlie differences in pain persistence or resolution remain poorly understood. This study focuses on sex-specific cellular phenotypes in the DRG following peripheral nerve injury and throughout the process of pain resolution. Our findings support the concept that pain resolution is an active biological process rather than a mere reversal of injury-induced changes. While the injury phase is dominated by an immune phenotype in both sexes, the pain resolution phase reveals distinct, sex-specific cell phenotypes and gene expression programs.

### In both sexes, pain resolution occurs without loss of neurons

The question of whether sensory neuron loss occurs after peripheral injury or under certain pain conditions is important (11, 13, 14, 16). In mice, it was shown that spared nerve injury (SNI) induces loss of small non-peptidergic neurons and reduces the DRG volume (14). In the rat after SNI, we could not find neuronal loss (16). Here, in rats of both sexes, no reduction or obvious change in abundance of sensory neurons was observed after CCI – neither after the injury nor during natural pain resolution. We also specifically examined potential subtype-specific loss of non-peptidergic neurons at both the cellular and molecular level. Neither the transcriptome data nor objective neuron counting showed evidence for loss of any neuron subtype. In accordance with older data, acquired in adult rats after sciatic nerve transection or crush (44), we assume that reduced numbers of non-peptidergic, IB4-positive, small diameter neurons after nerve injury may represent less immunoreactivity of the IB4 marker, rather than cell death. In humans, however, there is good evidence for disease-specific neuronal loss (11, 13). In humans suffering from diabetic painful neuropathy, DRG tissue shows substantial sensory neuron loss (11) and local formation of Nageotte nodules, which are local accumulations of satellite glia and non-myelinating Schwann cells, intertwined with sprouting sensory axons (12). In our rats, after CCI, such fundamental histological changes, like local accumulation of glial cells on cost of neuron abundance was not observed.

To our surprise, in males, but not females, the DRG tissue area harboring the sensory neurons was growing over time. We think that this growth in tissue size coincided with male body growth in this life span (7-12 weeks after birth). Consequently, during the experiment, neuronal density changed in males, but not females. Whether this is biologically relevant is not known, but the data underscore the need to carefully consider sex- and age-related factors in sensory neuron analyses. Notably, the overall number of sensory neurons seems to be higher in male than female rat DRG, at least in L4/L5.

### In both sexes, signaling mediators are massively upregulated after injury

CCI injury, in males and females, induces massive expression of signaling mediators such as *Npy*, *Vip*, *Il24*, *Gal*, and *Il6*, and macrophage-associated cytokine/chemokines such as *Csf1*, *Ccl2*, and *Il34* – all are still abundantly expressed in the resolution phase. For example, the most regulated neuropeptide was *Npy*, a factor well known for its function in vascularization (45). *Npy* was barely expressed in control DRG (∼3 TPM) but increased to1,300 TPM at both one and five weeks after CCI. This sustained and pronounced upregulation raises the question of whether *Npy*, or any of the other signaling peptides such as *Vip*, *Gal*, *Il6*, *Csf1*, and *Ccl2,* act just within the DRG microenvironment or exert broader systemic effects. In the DRG, NPY may act locally on vascularization or on DRG pericytes, which abundantly express the NPY receptor 1 (*Npy1r*, according to harmonized pain atlas (40)). It is also possible that NPY and other signaling mediators produced in the DRG exert effects beyond local circuits, contributing to long-distance DRG–body communication (46), although the underlying distribution mechanisms remain to be identified.

### Resolution-related transcriptomes are sex-specific

Pain resolution is not merely a reversal of the injury, but an active process marked by resolution-specific molecular and cellular phenotypes. In the injury phase, males and females show mostly overlapping transcriptional responses (Stephens et al., 2019). In our data, 58% (at log2FC ≥1) of all genes regulated during the injury phase were shared between both sexes. However, in the pain resolution phase, the intersecting gene set between males and females is reduced to ∼13% (log2FC ≥0.5). Notably, sex differences during this phase were most pronounced in immune-related processes, peptidase- and protein digestion activity, cell-cell junction assembly and signaling/excitability-related biological processes.

Differentially expressed genes (DEGs) after CCI were manifold and many of these are genes of unknown significance (GUS) in the context of pain. However, some of the sex-specific factors are well-known in pain research. The most significant sex-specific gene regulation pattern was found for the µ-opioid receptor *OPRM1*. This receptor has been shown to be associated with sex differences in antinociceptive responses (47), maintenance of neuropathic pain (48), and ethnicity-related effects in pain sensitivity (49). In humans and rodents, *OPRM1* is abundantly expressed in a subpopulation of peptidergic C-nociceptors, positive for *TRPV1*, *SCN10A* (Nav1.8), and *SCN11A* (Nav1.9) (50). In mice, conditional deletion of *Oprm1* specifically in *Trpv1*-positive cells, abolished morphine tolerance without affecting its antinociceptive effects (51). Women generally consume and receive lower opioid doses for acute and chronic pain, though it’s unclear if this reflects biological differences or prescribing patterns.

We also found sex-specific regulation for potassium ion channels – their expression was typically downregulated after injury (top candidates: *Kcnh7*, *Kcnb2*, *Kcns3*, and others). In females, *Kcnb2* (gene for Kv2.2) was downregulated after injury but re-expressed during resolution. In the DRG, Kv2 mRNA is widely expressed in many types of sensory neurons and is thought to limit neuronal excitability at rest (52). Dynamic regulation of its abundance, down after injury and up during resolution, may be one mechanism by which excitability is sex-specifically regulated in neuropathic pain (52).

### Sex-specific phenotype of macrophages at the interface between neurons and SGC

After peripheral nerve injury, sensory neurons, SGCs, as well as macrophages, are known to show massive transcriptional reprogramming and cell subtype diversification (36, 41, 53–56). Local macrophages in the DRG act sex-specifically and are key to initiation and persistence of neuropathic pain (36). Macrophages start to proliferate and self-renew locally, with cell phenotype differentiation occurring within hours (36, 55). In the DRG, many macrophages are found in close proximity to the sensory neuron-SGC unit. After injury, they strive for direct contact with sensory neurons and enter regions typically occupied by SGCs (37, 57), possibly to promote active pain resolution and nerve regeneration via mitochondria transfer (10, 55).

Our data suggest that the macrophages displace the SGCs from the neuron-SGC contact side after injury but retreat from this space during pain resolution. Concurrently, SGCs display sex-specific changes in their proximity to corresponding sensory neurons and in their cellular phenotype. In females, injury-induced GFAP expression decreased during pain resolution, whereas in males, GFAP expression remained elevated and even appeared on the contralateral side. The data raises the question of which active, regulated processes are needed to open and re-seal the neuron-SGC interface. Our transcriptome analysis suggests that tissue remodeling and regulation of cell contact, involving peptidases, protein turnover, and extracellular matrix, along with signaling and cell excitation regulation, are likely involved. It is probable that all three cell types, the neurons, the macrophages, and the glial cells, act coordinated. For intercellular communication, the transcriptome data provides multiple signaling candidates, including cytokines, chemokines, neuropeptides, signaling mediators, and cell-cell contact proteins, which are strongly upregulated after injury and remain expressed while pain resolution is ongoing.

In the past, research on sex as a biological factor of pain focused on sex-specific hormones, receptors, immune cells, and genetic factors (18, 28, 49, 58). Our data support the view that virtually the cell-specific gene expression landscape of DRG cells should be considered as biological variable when studying sex differences in pain processing.

### Strenghts and limitations of the study

Analysis of immunofluorescence microscopy images is a standard method to characterize cellular phenotypes in this field of research, but this approach is often applied to relatively small bioimage datasets with heuristic analysis. Moreover, applied analysis methods make it difficult to control for technical, anatomical, and analytical variability, potentially obscuring purely biological or sex-specific differences. We addressed this challenge on three levels: 1) We defined ongoing pain resolution à priori and experimentally. 2) We systematically analyzed whole tissue slices of individual DRGs in 2D, using computationally objective methods to evaluate all available image data. 3) We provide all bioimage data as an open-source dataset – comprising ∼7,000 tiles derived from ∼50,000 single images – serving as a reference for re-analysis or as potential training data. We note a risk that the dataset could be misused with AI to generate fabricated data. Although our 2D approach cannot provide definitive 3D neuron counts, it offers a valuable baseline for validating future tissue-clearing and quantitative labeling strategies aimed at fully characterizing sex-specific DRG neuron numbers across species.

On the single slice level, we found striking variability in the spatial distribution and abundance of labeled neurons, SGCs, and local macrophages. This variability implies that small datasets are unlikely to capture representative DRG regions, raising critical questions about the reliability of cell abundance analyses based on single DRG slices. Our data support the need for highly systematic whole-DRG analyses in 2D or 3D, with careful consideration of sex-specific cell phenotypes and à priori definition of sample sizes for each sex. We report the risk that given the multidimensional nature of our analyses, some statistically significant effects may represent random variation rather than true biological differences, when data categories build on low n-numbers (e.g. cell body area analysis in Fig. 2E).

We note the limitation that we failed to acquire complete ATF3 immunofluorescence data from female rats. For ethical reasons (3R, reduction), we did not repeat ATF3 labeling in new cohorts of female rats, as mRNA data were already available and showed comparable *Atf3* regulation between females and males. These considerations were also the basis for not including naïve animals and comparing the ipsi- to the contralateral side. We are aware that there is contralateral activity in DRGs (59) . However, using the contralateral side as a reference offers the advantage that smaller changes occurring bilaterally are excluded.

## Conclusion

Our study provides a framework for analyzing sex as a biological variable at the cellular level through deep analysis of large bioimage datasets. In the rat CCI model, this approach revealed sex-specific molecular and cellular phenotypes of sensory neurons, satellite glial cells and local macrophages, both after injury and during pain resolution. The data show that particularly the pain resolution phase is sex-specific, and marked by multi-cellular, multi-categorical biological processes. In summary, our study provides extensive resources for sex-specific resilience factors promoting pain resolution.

## RESOURCE AVAILABILITY

### Lead contact

Further information and requests for resources and reagents should be directed to and will be fulfilled by the Lead Contact, Robert Blum.

### Materials Availability

The study did not create new materials.

### Data and Code Availability

Raw and processed data were deposited within the Gene Expression Omnibus (GEO) repository (https://www.ncbi.nlm.nih.gov/geo) with an accession number (GSE278227). Custom Python scripts, deep learning models, and calculation scripts are available at GitHub (https://github.com/FSchlott/ratDRG_phenotyping_CCI_Sex_Resolution). The bioimage dataset is publicly available via the BioImage Archive (https://www.ebi.ac.uk/biostudies/bioimages/studies/S-BIAD1944; DOI: 10.6019/S-BIAD1944) and can be viewed with Streamlit (https://drg-pain-resolution.streamlit.app/).

## STAR METHODS

**table 2.** Key resources table

| REAGENT or RESOURCE | SOURCE | IDENTIFIER |
| --- | --- | --- |
| **Primary antibodies – dilution from stock** |  |  |
| ATF3 (polyclonal rabbit) | Sigma | #HPA001562<br>RRID: AB_1078233 |
| FABP7 (monoclonal rabbit, D8N3N) | Cell Signaling | #13347 |
| GFAP (polyclonal rabbit) | OriGene Technologies | #DP-014<br>RRID: AB_1001789 |
| GS (polyclonal mouse) | BD Transduction Laboratories | #610517<br>RRID: AB_397879 |
| Iba1 (polyclonal goat) | Fujifilm Wako Chemicals | #011-27991 |
| Isolectin B4, FITC conjugated | Sigma | #L2895 |
| NF-H (polyclonal chicken) | Millipore | #AB5539<br>RRID: AB_11212161 |
| **Secondary antibodies (all stored as 0.5 mg/ml)** |  |  |
| Alexa488; anti-chicken (Donkey) | Jackson Immuno Research | #703-545-155<br>RRID: AB_2340375 |
| Alexa488; anti-mouse (Donkey) | Invitrogen | #A-21202<br>RRID: AB_141607 |
| Alexa647; anti-chicken (Donkey) | Jackson Immuno Research | #703-605-155<br>RRID: AB_2340379 |
| Cy3; anti-rabbit (Donkey) | Jackson Immuno Research | #711-165-152<br>RRID: AB_2307443 |
| Cy5; anti-goat (Donkey) | Jackson Immuno Research | #705-175-147<br>RRID: AB_2340415 |
| **Chemicals, Peptides and Recombinant Proteins** |  |  |
| Aquapolymount | Polysciences Inc. | #18606 |
| Aqua | Braun | #0082479E |
| Aqua/Polymount | Polysciences | #18606 |
| DAPI | Sigma-Aldrich | #D9542 |
| dNTP Mix 10mM | Jena Bioscience | #NU-1006S |
| Donkey Serum | Sigma-Aldrich | #D9663 |
| Ethanol absolute | ChemSolute | #2246.1000 |
| Glycine | ITW Reagents | #131340 |
| Heparin-Natrium | Ratiopharm | PZN: 3029843 |
| Isofluran 1ml/ml | CP pharma |  |
| Nuclease-Free Water | Invitrogen /Ambion | #AM9932 |
| OCT Tissue Tec | Sakura | #4583 |
| Paraformaldehyde | Sigma-Aldrich | #P6148 |
| PBS | Sigma-Aldrich | #D8537 |
| Primer random p(dN)6 | Roche | #11034731001 |
| RNaseZap | Invitrogen / Ambion | #AM9782 |
| ROX Solution | Thermo Fisher | #R1371 |
| Sucrose | Sigma-Aldrich | #S7903 |
| Natriumethyl-mercurithiosalicilat (Thimerosal) | Roth | #6389.1 |
| Triton X100 | Sigma-Aldrich | #23.472 |
| Tween 20 | Roth | #9127.1 |
| **Critical Commercial Assays** |  |  |
| Arcturus <sup>TM</sup> PicoPure <sup>TM</sup> RNA Isolation Kit | Thermo Fisher / applied biosystems | #12204-01 |
| Luminaris HiGreen qPCR Master Mix | Thermo Fisher | #K0993 |
| RNasin® Plus Ribonuclease Inhibitor | Promega | #N2611 |
| RNeasy Mini Kit | Qiagen | #74104 |
| Superscript <sup>®</sup> III Reverse Transcriptase | Invitrogen | #18080-044 |
| **Experimental Models: organisms/strains** |  |  |
| Wistar rat | Janvier Labs, Saint Berthevin, France | Rj:Wi |
| **Software and Algorithms** |  |  |
| Anaconda Navigator | Anaconda Inc.<br>https://www.anaconda.com/products/individual-d |  |
| Cytoscape | https://cytoscape.org/ | (60) |
| Fiji Imaging Software | https://fiji.sc/ | (61) |
| G-Power |  | (62) |
| Metascape | https://metascape.org/ | (63) |
| Python 3.8 | Python Software Foundation<br>https://www.python.org/ |  |
| QuPath (v0.3.1) | https://qupath.github.io/ | (64) |
| Streamlit | Streamlit Inc.<br>https://streamlit.io/ |  |
| Visual Studio Code | https://code.visualstudio.com/ |  |
| Zen Imaging Software Version 2.6 | Carl Zeiss Microscopy GmbH |  |
| **Technical Devices / Other** |  |  |
| Axiolmager2 | Zeiss |  |
| IX81 microscope | Olympus |  |
| MC3050 S Kryostat | Leica |  |
| MM 400 (Bead Mill) | Retsch |  |
| NanoDrop™ OneC Microvolume UV-Vis Spectrophotometer | Thermo Fisher |  |
| StepOnePlus | Applied Biosciences |  |
| ThermoMixer C | Eppendorf |  |
| T3 Thermocycler | Biometra |  |

## EXPERIMENTAL MODEL AND SUBJECT DETAILS

### Experimental strategy (detailed version)

We modeled pain resolution by performing chronic constriction injury (CCI) of the sciatic nerve in adult male and female rats (Figure 1A). À priori, we defined the resolution stage as a 50% reduction in mechanical hypersensitivity, measured by von Frey testing (65), to capture a transitional state before full recovery (32). Consistent with previous work from our group, peak mechanical hypersensitivity occurred within two weeks post-CCI, and the 50% recovery threshold was consistently reached by five weeks in both sexes, with no significant sex differences (Figure 1A and Figure S1).

For cellular phenotyping, single DRGs from lumbar segments L4 and L5 – known to exhibit strong plasticity responses to traumatic sciatic nerve injury (16) – were harvested and processed for systematic bioimaging using tile and confocal microscopy. These analyses addressed neuronal abundance, satellite glial cell (SGC) and macrophage (Mph) phenotypes, and structural changes in DRG units (sensory neurons, SGCs, and local macrophages) (Figure 1B). mRNA sequencing from single DRGs generated molecular profiles that complemented the imaging data and revealed gene expression plasticity during pain resolution in both sexes (Figure 1B). Statistical analyses assessed injury-, resolution-, and sex-specific phenotypes using ipsilateral and contralateral DRGs collected either one-week post-CCI (injury stage) or five weeks post-CCI (resolution stage, 50% recovery from mechanical hypersensitivity) (Figures 1C–E). Here, the term lateralization refers to phenotypes emerging on the contralateral side during the course of pain resolution (Figure 1C).

For bioimage analysis, we used a deep learning (DL)-based image feature segmentation strategy (34) implemented in the pipeline deepflash2 (33), specifically optimized for DRG tissue (13, 16) (Figure 1F).

For cellular phenotyping, we performed serial sectioning and immunolabeling of entire L4 and L5 DRG, across 10 slides per sample (Figure 1F). After immunofluorescence microscopy, corresponding immunolabels were independently annotated by three experts, a computational consensus was created, and DL model ensembles were trained (Figure 1F and Figure S2). Images were used to compute segmentation masks for the following labels: neurofilament (NF) for neuronal cell bodies, IB4 for non-peptidergic neurons, ATF3 as neuronal injury marker, FABP7 for SGC, and glutamine synthetase as well as GFAP as plasticity markers for SGC (Figure S3) (13, 16).

Neurons and SGCs were segmented using DL model ensembles, achieving dice similarity scores ≥0.85 on both validation and test image sets (Tables S1, S2, S3). Tissue area and macrophages, identified with Iba1 (13), were segmented using advanced thresholding approaches (Figure S3).

This way, we analyzed a total of approximately 6,914 tile microscopy images, each consisting of 4 – 10 stitched sub-images covering entire DRG cross sections. The 6-16 slices per DRG contained on average 1,000-3,000 neurons in a tissue area of 7 - 13 mm^2^ per DRG for each staining (Figure S4). High resolution spatial analysis of the neuron-SGC and macrophage units was done on 576 confocal images.

### Rats

All experiments were in accordance with the guidelines set by the European Union and approved by our institutional Animal Care, the Utilization Committee and local authorities (Regierung Unterfranken, RUF55.2.2-2532-2-612 and -2-1752). Experiments were in accordance with the ARRIVE guidelines.

Male and female Wistar rats were delivered with an age of 5 weeks (weight ∼150 – 175 g) were housed with environmental enrichments in groups of 3 to 5 rats per cage in the local animal facility (ZEMM, University of Würzburg, Germany). Before the start of the experiments, rats were habituated to the new housing for a week. Rats housed at a 12h/12h light/dark circadian cycle at 21-25°C and 45-55°C humidity, with food and water *ad libitum*. Rats were pathogen-free. All behavioral experiments were performed at an age of 6–12 weeks during the subjective day-phase of the animals and were randomly allocated to experimental groups. The minimal group size was calculated by à priori power analysis with G-Power 3.1 (difference between two independent means, group comparison) based on the effect size of the behavioral results, which was close to 1.8 (α-error 0.05, power 0.8). Rats were perfused at an age of 12 weeks and showed a mean weight of ∼440 g (males) or ∼260 g (females).

## METHOD DETAILS

### Chronic constriction injury

Surgery was performed under deep isoflurane anesthesia (3% isoflurane, O_2_). An incision was placed above the upper thigh, and the muscle was bluntly opened to expose the sciatic nerve. The nerve was freed of connective tissue, and four loose silk ligatures were placed around the nerve, spaced 1 mm apart. The silk thread was soaked in 0.9% saline before use. The wound was closed using a non-absorbable surgical suture (Optilene 3/0, #3090435 Braun, Germany). After wound closure, the skin was disinfected with Octeniderm solution (Schülke and Mayr, Germany). After CCI, the animalś wellbeing was assessed daily, using appropriate score sheets.

### Behavioral Testing

To examine pain behavior after CCI, the von Frey up-down method was used (65). As effect variable, the 50% withdrawal threshold was calculated. Animals were habituated to the handling and von Frey setup daily for 30 minutes each one week prior to the CCI. Behavioral tests were done before the surgery (d0, baseline) and each week until the average 50% withdrawal threshold had increased more than 50% back to the baseline value compared to the value of highest hypersensitivity. For the von Frey test, animals were placed on a mesh platform in boxes and acclimatized for at least 5 min before behavioral testing. Testing started with a 4 g filament. The animal tested had to sit still on all four paws during the application of the stimulus. The filament was applied to the plantar surface of the hind paw for 1-1.5 s until the filament buckled. Withdrawal of the paw, flinching, and licking were counted as a positive behavioral response. The results were recorded and the 50% withdrawal threshold was calculated as described (66).

### Perfusion fixation for immunostaining

Animals were anesthetized with 4% isoflurane (in oxygen) and a mixture of ketamine-xylazine (90mg/kg body weight (BW); 10mg/kg BW, i.p.). The abdominal wall and ribcage were opened to expose the heart. A needle was injected into the left ventricle, and the right atrium was opened for perfusion. Animals were first rinsed with 0.4% heparin in PBS (approx. 100ml) and subsequently fixed with a 4% paraformaldehyde (PFA) solution (approx. 200ml per animal). Inner organs and muscles were removed to access the vertebral column, which was cut out above and below the lumbar vertebrae L4 and L6. The spinal column was opened on the ventral and dorsal sides and the DRGs of L4-L5 were collected from the ipsilateral and contralateral sides in ice-cold PBS. DRGs were post-fixed in 4% PFA for another 30 min. DRGs were shortly rinsed in PBS and free formaldehyde was quenched for 1 h with 100 mM glycine (pH 7.4 with Tris-base). Afterwards, all samples were washed three times in PBS. Dehydration was performed in 30% sucrose overnight at 4°C. The DRG were embedded in Tissue-Tek O.C.T. using dry ice and were stored at −80°C.

### Cryosectioning and immunostaining

Both L4 and L5 DRGs were processed. Each DRG was cut at −24°C in 16 µm thick slices using a Leica CM3050 S Cryostat. The slices were collected serially on 10 object slides, with every 10th slice per slide, so that each object slide represented an entire DRG (Figure 1F).

For immunostaining, free formaldehyde was again quenched in 100 mM glycine (pH 7.4 with Tris-base) for 30 min. Slides were washed twice with PBS for 5 min and afterwards incubated in blocking solution (10% donkey serum, 0.3% Triton X100 and 0.1% Tween20 in PBS) for one hour. Primary antibodies were applied at 4°C, overnight. After washing five times with wash buffer (0.1% Triton X100 and 0.1% Tween 20 in PBS) for two min each, secondary antibodies were added in blocking solution and incubated at room temperature for 1 h. The slides were washed three times in washing buffer for 2 min and then once more in PBS for 5 min. DAPI (2 mg/ml stock solution, freshly diluted 1/5,000 in PBS) was applied for 10 min. Slides were washed three times in PBS and for 5 min each and shortly desalted with water. Aqua/Polymount was used as mounting medium. The slides were sealed with nail varnish and stored at 4°C.

### Tile microscopy (epifluorescence)

Tile microscopy was performed using an AxioImager2 microscope (Zeiss) using a 20× Plan-Apochromat objective (0.8 N.A). Images with 14-bit were acquired using an Axiocam 506 mono camera (Zeiss) with a binning of 4×4 at a pixel resolution of 0.908 µm/pixel. As a light source, the LED excitation wavelengths (in nm) 385, 475, 555, and 630 of the Colibri.2 LED light source (Zeiss) were used together with corresponding multiband filter sets. Whole DRG were imaged with the tile scan function of the Zen Imaging Software (Version 2.6). For each DRG, 6 – 12 x,y (2D)-images were stitched together. LED intensities were adjusted to elicit the strongest signal in the first third of the 14-bit scale and each staining was imaged with the same settings. Raw images were stored in the .czi format.

### Confocal microscopy

High-resolution confocal images with were acquired at an inverted Olympus IX81 microscope equipped with an Olympus FV1000 confocal laser scanning system. A FVD10 SPD spectral detector and diode lasers of 405, 473, 559, and 635 nm were used in combination with an Olympus UPLSAPO60× (oil, NA: 1.35) objective. The pinhole was set to represent one airy disc. Confocal images (x,y-z; 12-bit, 1024×1024 pixel) were taken throughout the depth of the tissue with approximately 20 to 30 planes (400 nm step size). Raw images were stored in the .oib format.

### Deep learning-based bioimage analysis

An overview of the image analysis process is given in Figure 1, S2 and S3. Czi-files that were acquired at the AxioImager2 microscope (Zeiss) were split into single channels and converted to tiff-file images using the tifffile package in Python. For confocal images, a maximum projection of the middle five planes was used, computed with oiffile and saved as tiff files. Due to higher background values in the staining of males, 5 weeks, image LUT settings had to be adjusted to reduce the background and apply the otsu-threshold. LUT settings were cut between values of 250 and 2500.

In the next step, binary segmentations were generated for each image with the use of deepflash2 (33) as demonstrated in (16) or, in case of Iba1 staining using a thresholding method. For the staining of NF and GS/GFAP, preexisting models were used (16). The preexisting NF model was trained and refined by adding 10 images from the female cohort to the NF dataset. For the staining of ATF3 and IB4, as well as confocal images of FAPB7 and NF staining, new models for segmentation of ATF^+^ neurons, AFT3^+/-^ neurons, IB4^+^ neurons, FABP7^+^ SGCs, and NF^+^ neurons were generated (Supplementary Table 1-3).

For creating a new model, 15 exemplary images were annotated by three blinded experts in QuPath (64) for each staining. The annotations were exported as JSON-files and converted to binary segmentations. The segmentations and corresponding images of each annotator were loaded onto the deepflash2 GUI and a ground truth estimation was performed with the STAPLE algorithm to generate one consensus segmentation from all three annotators. 80% (twelve images) of the ground truth estimation were then used for the Ensemble Training. Three models were trained using the default settings (ATF3) or adjusted to a decreased learning rate (0.0003, IB4) to create an ensemble model. Of the twelve images, three were used for validation during training. Afterwards, the three remaining, yet by the model unseen images (20%), were used for quality control to evaluate the performance of the model ensemble.

The Iba1 staining, which indicates macrophages, was not suited to manual annotations and a deep learning approach because of the intricate processes of the macrophages. Instead, a thresholding approach was used. To first reduce the background, images were sharpened using the “unsharp_mask” function in the Python library “skimage.filters” for tile images; Confocal images were sharpened using the “gaussian” function. Then, a segmentation was created based on the Otsu’s method (skimage.filters.otsu_threshold) and the noise was reduced with a median filter.

The deep learning models and Iba1-thresholding method were applied to all acquired images. The quality of the predicted segmentations was revised by screening each image and overlay in an application in streamlit.io (publicly available data viewer at https://drg-pain-resolution.streamlit.app/). A few images with deficient segmentations due to staining problems were removed prior to further analysis.

Analysis of the images and corresponding segmentations was done in Python3. The mean values then were accessed for statistics and plotting in Jupyter notebook (Anaconda). Files are openly available on GitHub (https://github.com/FSchlott/ratDRG_phenotyping_CCI_Sex_Resolution).

### RNA isolation

Animals were euthanized with CO_2_. The L4 and L5 DRG were collected from both the ipsilateral and the contralateral side. All samples were snap-frozen on liquid nitrogen immediately after collection. Samples were stored at −80°C until further use.

RNA was extracted and purified using the Arcturus PicoPure RNA Isolation Kit. The frozen tissue samples were emersed in 100 µl extraction buffer. Cell lysis was performed using the Retsch MM400 bead mill with one zirconium oxide 5 mm grinding ball at 30 Hz for 2 min at room temperature. Tubes were shortly centrifuged and incubated at 42°C for 30 min in a ThermoMixer. RNA isolation was performed according to the manufacturer’s instructions. After the second application of wash buffer 2, the columns were dried at max speed for three minutes. Samples were eluted in 15 µl elution buffer. RNA concentration was measured at the NanoDrop OneC and stored at −20°C until further use. We received about 2.2 µg total RNA per DRG for each group.

### Reverse Transcription and qPCR

Reverse Transcription of 500ng of L5-DRG RNA was carried with Superscript III Reverse Transcriptase Kit. For cDNA synthesis, RNA was incubated for 5 min at 65°C followed by 5 min at 4°C with 20pmol dNTPs, random hexamer primers, and RNasin Plus Ribonuclease Inhibitor. First Strand Buffer and DTT (ad 5mM) were added, and the mixture was incubated at 37°C for 4 min. For reverse transcription 1 µl Superscript III (200 units) was added to the mix to reach a final volume of 20 µl. The reaction was incubated at 37°C for 2 h and stopped at 70°C for 15 min. Samples were stored at −20°C until further use. Before performing the qPCR, RT samples were diluted 1/5 and the qPCR was performed with the Luminaris HiGreen qPCR Master Mix with SYBR Green as the detection agent. ROX solution was added to a concentration of 600 nM as internal fluorescence reference. 2 µl of cDNA (representing 10 ng RNA) and 6 pmol of forward and reverse primers were used to a final volume of 20 µl. Thermal cycling was performed in the StepOnePlus Cycler in a two-step protocol, as suggested by the Luminaris HiGreen qPCR Master Mix protocol (UDG pre-treatment at 50°C for 2 min and initial denaturation at 95°C for 10 min. 40 cycles of 15 s denaturation at 95°C and 60 s annealing and extension were run. A melting-curve was generated after each PCR run (15 s 95°C, 1 min 60°C, 15s 95°C).

Reverse transcription–qPCR analysis was performed for the housekeeping gene *Gapdh* and selected injury-related (*Gfap*, *Abca1*, *ApoE*) and inflammation-related genes (*Il1b, Il6, Tnfα, Tgfβ, Nox2, Il10, Igf1*) using one cohort (males, 1 week, ipsi- vs. contralateral) (Figure S10). These data confirmed comparable RNA input across samples and verified that CCI significantly increased expression of cell-plasticity markers (e.g., *Gfap, Abca1, Nox2*) and the low-abundance inflammatory marker *Il6*.

After reverse transcription, we found virtually no difference in abundance (p = 0.559) for the housekeeping gene *Gapdh*, contralateral (ct = 18.18 ± 0.14 SD, n = 6, corresponding to ∼150,000 copies of *Gapdh* per 2.5 ng input RNA; ipsilateral (18.09 ± 0.31; n = 7)). Following RNA-Seq and transcript per million (TPM) normalization (supplementary data 1), we found in the same samples ∼1,570 transcripts per million (TPM) of *Gapdh* contralateral and ∼1,690 TPM ipsilateral. For the low abundant *Il6*, we sequenced 0.14 TPM contralateral and 8 TPM ipsilateral. The data confirmed comparable RNA quality across samples and likely representation of low-abundant transcripts, such as *Il6*, within the bulk RNA data.

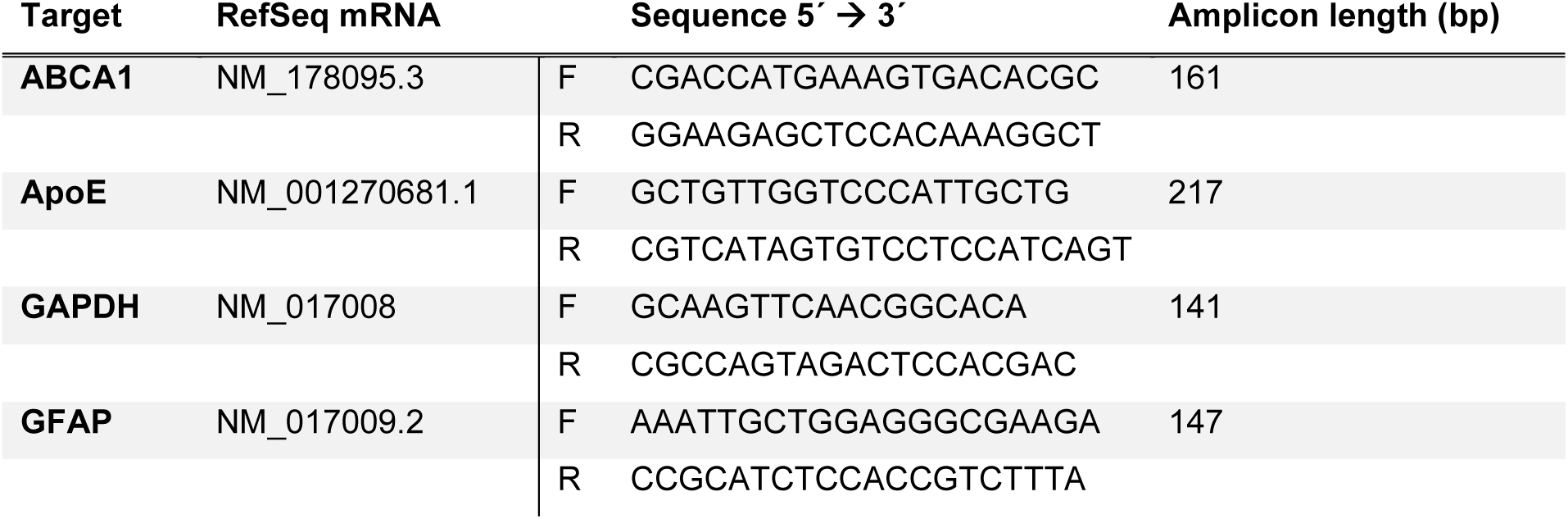
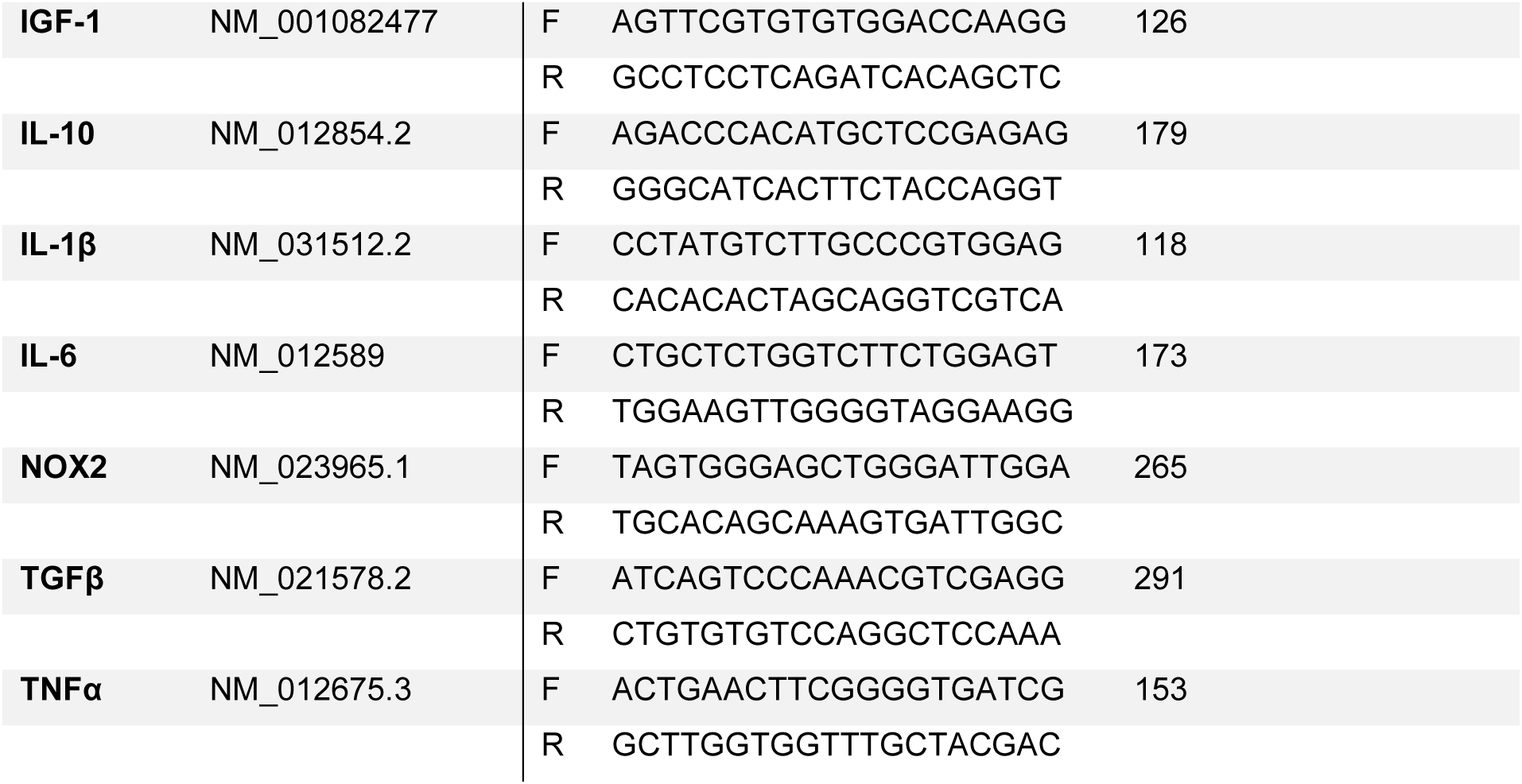

### RNA sequencing

RNA quality was checked using a 2100 Bioanalyzer with the RNA 6000 Nano kit (Agilent Technologies). The RIN for all samples was ≥ 7,5. DNA libraries suitable for sequencing were prepared from 300 ng of total RNA with oligo-dT capture beads for poly-A-mRNA enrichment using the TruSeq Stranded mRNA Library Preparation Kit (Illumina) according to manufacturer’s instructions (1/2 volume). After 15 cycles of PCR amplification, the size distribution of the barcoded DNA libraries was estimated ∼295 bp by electrophoresis on Agilent DNA 1000 Bioanalyzer microfluidic chips. Sequencing of pooled libraries was performed at 20-40 million reads/sample in single-end mode. Male/1-week libraries were sequenced with 76 nt read length on the NextSeq 500 platform (Illumina) while male/24h, male/5-week and all female libraries were sequenced at 101 nt read length on the NextSeq 2000 platform (Illumina). Demultiplexed FASTQ files were generated using bcl2fastq2 v2.20.0.422 (Illumina) for male/1-week and male/5-week data, and bcl-convert v4.0.3 (Illumina) for the remaining data.

To ensure high sequence quality, Illumina reads were quality- and adapter-trimmed using Cutadapt (67) version 2.5 using a cutoff Phred score of 20 in NextSeq mode, and reads without any remaining bases were discarded (parameters: --nextseq-trim=20 -m 1 -a AGATCGGAAGAGCACACGTCTGAACTCCAGTCAC). For comparison with male/1-week samples, male/5-week data was initially trimmed by 25 bases from read 3’ ends (parameters: -u −25 --nextseq-trim=20 -m 1 -a AGATCGGAAGAGCACACGTCTGAACTCCAGTCAC) but processed normally for comparison with female/5-week data.

Processed reads were subsequently mapped to the rat genome (GCF_015227675.2/mRatBN7.2) using STAR (68) v2.7.2b with default parameters but including transcript annotations from RefSeq annotation version 108 for mRatBN7.2. The mRatBN7.2 v108 annotation was also used to generate read counts on exon level summarized for each gene via featureCounts v1.6.4 from the Subread package (69).

## QUANTIFICATION AND STATISTICAL ANALYSIS

### Transcriptome analysis

Multi-mapping and multi-overlapping reads were counted strand-specific and reversely stranded with a fractional count for each alignment and overlapping feature (parameters: -s 2 -t exon -M - O --fraction). The count output was used to identify differentially expressed genes using DESeq2 (70) version 1.24.0. Read counts were normalized using DESeq2, and fold-change shrinkage was applied by setting the parameter “betaPrior=TRUE”. Three independent DESeq2 analyses were conducted with data for (i) male/1-week and male/5-weeks (76 nt), (ii) male/24h and females as well as (iii) male/5-weeks (101 nt) and female/5-weeks.

For transcriptome analysis, DESeq2 data were used. Volcano plots and heatmaps were generated with Python3. Gene expression heatmaps of the top regulated genes show the mean for each condition of the normalized values, filtered for protein coding genes only. All significantly regulated genes (padj > 0.01) above a Log2 fold change cutoff were taken into account for the subsequent pathway analysis on metascape.org. This cutoff was at a change of Log2FC ±1 for the injury transcriptome (CL vs. IL side) and was lowered to Log2FC ±0.5 for the resolution transcriptome (1W vs. 5W) to achieve a sufficient number of regulated genes for enrichment analysis. Protein-protein interaction networks were generated with Metascape (63) and visual adjustments were made with Cytoscape (60).

### Calculations in bioimage analysis

Calculation details are available on GitHub: (https://github.com/FSchlott/ratDRG_phenotyping_CCI_Sex_Resolution).

*Injury response of neurons after chronic constriction injury across neuronal subtypes.:*

Total numbers displayed for the tissue area and neuron number are the sum of the values per slice for tissue area and neuron numbers, respectively. For neurons, any segmentation labels smaller than the smallest annotated size (122.1µm^2^) were excluded from the analysis. For the histograms (Figure 2E, Figure 3E), the number of NF, IB4, and ATF3 positive neurons were divided into bins of 200 µm^2^. The total number of NF neurons was defined as 100% and the fraction of neurons for each bin was calculated. Scatterplots (Figure 2G, Figure 3F) show the relation between neuron number and tissue area and represent one slice per datapoint. For comparing the distribution of IB4 per NF and ATF3 for each subtype, the percentage of an overlap between each staining was calculated.

*Activation of satellite glial cells and macrophages after chronic constriction injury:*

In order to focus on the neuron-near area (NNA), the NF segmentation was dilated with the “binary_dilation” function from the Python library “skimage.morphology” and only the Iba1 and GFAP segmentations in this NNA were quantified. The positive area was calculated as the sum of positive pixels within the NNA and the number of labels was determined (skimage.measure.labels).

#### The neuron-SGC units and macrophages

To measure the contact or close proximity of SGC and Mph to neurons, we introduced a parameter for the neuron segmentation outline (NO) and its overlap with the respective Iba1 and GS / Fabp7 segmentations. For this, we iterated over each neuron and set the border of each NF segmentation as neuron border. To include close proximity, the Iba1 / GS / Fabp7 masks were dilated by 1px before calculating the percentage of overlap with the NO. A mean per neuron was then calculated per slice and proceeded as before. To quantify the number of neurons that were, in any capacity touched by SGC or Mph, we quantified neurons positive for any overlap in percent to the total neuron number.

### Statistical analysis

As shown in Figure 1, we à priori set the following hypotheses to examine injury-, resolution-, and sex-dependent effects in our DRG dataset: We tested for a) injury related effects (*H_0_: there is no difference between the ipsilateral and contralateral side at 1W and 5W after injury*), b) pain resolution related effects (*H_0_: there is no change on the ipsilateral side between 1 week and 5 weeks post-injury*), and c) lateralization of effects to the contralateral side over time (*H_0_: there is no difference between the contralateral sides at 1W and 5W after injury*), analyzed separately for male and female animals. Next, we tested for sex differences d) at baseline (*H_0_: there is no difference between male and female animals on the contralateral side*) and e) after injury (*H_0_: there is no difference between male and female animals on the ipsilateral side*), both for 1W and 5W. Lastly, in case of the ATF3 staining, we did ATF3 stratified comparisons (*H_0_: there is no difference between ATF3 positive and negative neurons on the ipsilateral side* and *H_0_: there is no difference between 1W and 5W within ATF3 positive and negative subpopulations on the ipsilateral side*). In all analyses, individual DRGs were treated as independent data points, based on the evidence that inter-DRG variability is comparable to inter-animal variability (16).

#### Statistical Workflow

All statistical analyses were performed using Python 3.12 and standard packages including “pandas”, “scipy”, “statsmodels”, and “numpy”. Each hypothesis was tested using a two-group comparison framework based on the following pipeline: First, we tested the data for the assumption of normality using the Shapiro-Wilk test. For unpaired comparisons, homogeneity of variances was tested using Levene’s test. Based on the results, for paired comparisons between the ipsi- and contralateral sides (samples from the same animals), a paired t-test (if normality was met) or a Wilcoxon signed-rank test (if not) was used. For unpaired samples, either an independent *t*-test (with equal variance) or a Mann–Whitney U test (without equal variance) was used. For each hypothesis group, p-values were corrected for multiple comparisons using the Bonferroni method. The significance level was set as p(adj) < 0.05, and is indicated in figures as (*p ≤0.05; **p ≤0.01; ***p ≤0.001).

## SUPPLEMENTAL INFORMATION

Supplemental information includes 3 supplementary tables, 11 supplementary figures, 12 RNA-Seq data files.

## ACKNOWLEDGEMENTS

We thank Niels Köhler, Martina Wilhelm, Ida Kanzerani, and Mariam Sobhy Attala for help with cell annotation.

This work was funded by the Deutsche Forschungsgemeinschaft (DFG, German Research Foundation), project ID: 426503586, KFO5001 ResolvePAIN (to A.B., H.L.R., and R.B.), the Evangelisches Studienwerk Villigst (to A.S.); and the Interdisciplinary Centre for Clinical Research Würzburg (IZKF) project Z-6 (to T.G.).

## AUTHOR CONTRIBUTIONS

Conceptualization, F.S., H.L.R. A.S., and R.B.; Methodology, F.S., A.S., T.B., T. G., and R.B.; Formal Analysis: F.S., T.B., T.G., A.S.; Investigation, F.S., B.H., A.S., R.B.; Data Curation, F.S.; Writing – Original Draft, F.S., A.S., and R.B.; Writing –Review & Editing, F.S., T.B., T.G., A.B., H.L.R., A.S., and R.B.; Supervision, A.B., R.B.; Project Administration, R.B.; Funding Acquisition, A.S., A.B., H.L.R., R.B.

## REFERENCES

1. Scholz J, Finnerup NB, Attal N, Aziz Q, Baron R, Bennett MI, et al. The IASP classification of chronic pain for ICD-11: chronic neuropathic pain. Pain. 2019;160(1):53–9.

2. Soliman N, Moisset X, Ferraro MC, de Andrade DC, Baron R, Belton J, et al. Pharmacotherapy and non-invasive neuromodulation for neuropathic pain: a systematic review and meta-analysis. Lancet Neurol. 2025;24(5):413–28.

3. van Hecke O, Austin SK, Khan RA, Smith BH, Torrance N. Neuropathic pain in the general population: a systematic review of epidemiological studies. Pain. 2014;155(4):654–62.

4. Cohen SP, Vase L, Hooten WM. Chronic pain: an update on burden, best practices, and new advances. Lancet. 2021;397(10289):2082–97.

5. Sandy-Hindmarch O, Bennett DL, Wiberg A, Furniss D, Baskozos G, Schmid AB. Systemic inflammatory markers in neuropathic pain, nerve injury, and recovery. Pain. 2022;163(3):526–37.

6. Hakim S, Jain A, Woolf CJ. Immune drivers of pain resolution and protection. Nat Immunol. 2024;25(12):2200–8.

7. Gangadharan V, Zheng H, Taberner FJ, Landry J, Nees TA, Pistolic J, et al. Neuropathic pain caused by miswiring and abnormal end organ targeting. Nature. 2022;606(7912):137–45.

8. Renthal W, Tochitsky I, Yang L, Cheng YC, Li E, Kawaguchi R, et al. Transcriptional Reprogramming of Distinct Peripheral Sensory Neuron Subtypes after Axonal Injury. Neuron. 2020;108(1):128–44 e9.

9. Scheib J, Höke A. Advances in peripheral nerve regeneration. Nature reviews Neurology. 2013;9(12):668–76.

10. van der Vlist M, Raoof R, Willemen H, Prado J, Versteeg S, Martin Gil C, et al. Macrophages transfer mitochondria to sensory neurons to resolve inflammatory pain. Neuron. 2022;110(4):613–26 e9.

11. Hall BE, Macdonald E, Cassidy M, Yun S, Sapio MR, Ray P, et al. Transcriptomic analysis of human sensory neurons in painful diabetic neuropathy reveals inflammation and neuronal loss. Scientific reports. 2022;12(1):4729.

12. Shiers SI, Mazhar K, Wangzhou A, Haberberger R, Lesnak JB, Sankaranarayanan I, et al. Nageotte nodules in human DRG reveal neurodegeneration in painful diabetic neuropathy. bioRxiv. 2024.

13. Sodmann A, Degenbeck J, Aue A, Schindehutte M, Schlott F, Arampatzi P, et al. Human dorsal root ganglia are either preserved or completely lost after deafferentation by brachial plexus injury. British journal of anaesthesia. 2024;133(6):1250–62.

14. Cooper AH, Barry AM, Chrysostomidou P, Lolignier R, Wang J, Redondo Canales M, et al. Peripheral nerve injury results in a biased loss of sensory neuron subpopulations. Pain. 2024;165(12):2863–76.

15. Kuo LT, Simpson A, Schanzer A, Tse J, An SF, Scaravilli F, et al. Effects of systemically administered NT-3 on sensory neuron loss and nestin expression following axotomy. J Comp Neurol. 2005;482(4):320–32.

16. Schulte A, Lohner H, Degenbeck J, Segebarth D, Rittner HL, Blum R, et al. Unbiased analysis of the dorsal root ganglion after peripheral nerve injury: no neuronal loss, no gliosis, but satellite glial cell plasticity. Pain. 2023;164(4):728–40.

17. Smith AF, Plumb AN, Berardi G, Sluka KA. Sex differences in the transition to chronic pain. The Journal of clinical investigation. 2025;135(11).

18. Mogil JSP-Z, E.M.; Roche, M.; Vincent, K.;. Fact sheets: Biological mechanisms underlying sex differences in pain 2024 [Available from: https://www.iasp-pain.org/resources/fact-sheets/biological-mechanisms-underlying-sex-differences-in-pain/.

19. Mogil JS, Parisien M, Esfahani SJ, Diatchenko L. Sex differences in mechanisms of pain hypersensitivity. Neurosci Biobehav Rev. 2024;163:105749.

20. Mogil JS. Qualitative sex differences in pain processing: emerging evidence of a biased literature. Nature reviews Neuroscience. 2020;21(7):353–65.

21. Collaborators GBDLBP. Global, regional, and national burden of low back pain, 1990-2020, its attributable risk factors, and projections to 2050: a systematic analysis of the Global Burden of Disease Study 2021. Lancet Rheumatol. 2023;5(6):e316–e29.

22. Mogil JS. Sex differences in pain and pain inhibition: multiple explanations of a controversial phenomenon. Nature reviews Neuroscience. 2012;13(12):859–66.

23. Ghazisaeidi S, Muley MM, Salter MW. Neuropathic Pain: Mechanisms, Sex Differences, and Potential Therapies for a Global Problem. Annu Rev Pharmacol Toxicol. 2023;63:565–83.

24. Presto P, Mazzitelli M, Junell R, Griffin Z, Neugebauer V. Sex differences in pain along the neuraxis. Neuropharmacology. 2022;210:109030.

25. Gulati M, Dursun E, Vincent K, Watt FE. The influence of sex hormones on musculoskeletal pain and osteoarthritis. Lancet Rheumatol. 2023;5(4):e225–e38.

26. Stratton H, Lee G, Dolatyari M, Ghetti A, Cotta T, Mitchell S, et al. Nociceptors are functionally male or female: from mouse to monkey to man. Brain. 2024;147(12):4280–91.

27. Avona A, Burgos-Vega C, Burton MD, Akopian AN, Price TJ, Dussor G. Dural Calcitonin Gene-Related Peptide Produces Female-Specific Responses in Rodent Migraine Models. The Journal of neuroscience : the official journal of the Society for Neuroscience. 2019;39(22):4323–31.

28. Fan CY, McAllister BB, Stokes-Heck S, Harding EK, Pereira de Vasconcelos A, Mah LK, et al. Divergent sex-specific pannexin-1 mechanisms in microglia and T cells underlie neuropathic pain. Neuron. 2025;113(6):896–911 e9.

29. Sorge RE, Mapplebeck JC, Rosen S, Beggs S, Taves S, Alexander JK, et al. Different immune cells mediate mechanical pain hypersensitivity in male and female mice. Nat Neurosci. 2015;18(8):1081–3.

30. Millecamps M, Sotocinal SG, Austin JS, Stone LS, Mogil JS. Sex-specific effects of neuropathic pain on long-term pain behavior and mortality in mice. Pain. 2023;164(3):577–86.

31. Farrar JT, Young JP, Jr., LaMoreaux L, Werth JL, Poole MR. Clinical importance of changes in chronic pain intensity measured on an 11-point numerical pain rating scale. Pain. 2001;94(2):149–58.

32. Hartmannsberger B, Ben-Kraiem A, Kramer S, Guidolin C, Kazerani I, Doppler K, et al. TAM receptors mediate the Fpr2-driven pain resolution and fibrinolysis after nerve injury. Acta Neuropathol. 2024;149(1):1.

33. Griebel M, Segebarth D, Stein N, Schukraft N, Tovote P, Blum R, et al. Deep learning-enabled segmentation of ambiguous bioimages with deepflash2. Nature communications. 2023;14(1):1679.

34. Segebarth D, Griebel M, Stein N, von Collenberg CR, Martin C, Fiedler D, et al. On the objectivity, reliability, and validity of deep learning enabled bioimage analyses. eLife. 2020;9.

35. Pinto LG, Souza GR, Kusuda R, Lopes AH, Sant’Anna MB, Cunha FQ, et al. Non-Peptidergic Nociceptive Neurons Are Essential for Mechanical Inflammatory Hypersensitivity in Mice. Mol Neurobiol. 2019;56(8):5715–28.

36. Yu X, Liu H, Hamel KA, Morvan MG, Yu S, Leff J, et al. Dorsal root ganglion macrophages contribute to both the initiation and persistence of neuropathic pain. Nature communications. 2020;11(1):264.

37. Iwai H, Ataka K, Suzuki H, Dhar A, Kuramoto E, Yamanaka A, et al. Tissue-resident M2 macrophages directly contact primary sensory neurons in the sensory ganglia after nerve injury. Journal of neuroinflammation. 2021;18(1):227.

38. Vega-Avelaira D, Géranton SM, Fitzgerald M. Differential regulation of immune responses and macrophage/neuron interactions in the dorsal root ganglion in young and adult rats following nerve injury. Molecular pain. 2009;5:70.

39. Avraham O, Deng PY, Jones S, Kuruvilla R, Semenkovich CF, Klyachko VA, et al. Satellite glial cells promote regenerative growth in sensory neurons. Nature communications. 2020;11(1):4891.

40. Bhuiyan SA, Xu M, Yang L, Semizoglou E, Bhatia P, Pantaleo KI, et al. Harmonized cross-species cell atlases of trigeminal and dorsal root ganglia. Sci Adv. 2024;10(25):eadj9173.

41. Jager SE, Pallesen LT, Richner M, Harley P, Hore Z, McMahon S, et al. Changes in the transcriptional fingerprint of satellite glial cells following peripheral nerve injury. Glia. 2020;68(7):1375–95.

42. Xu L, Chen Z, Li X, Xu H, Zhang Y, Yang W, et al. Integrated analyses reveal evolutionarily conserved and specific injury response genes in dorsal root ganglion. Sci Data. 2022;9(1):666.

43. Avraham O, Hadas Y, Vald L, Zisman S, Schejter A, Visel A, et al. Transcriptional control of axonal guidance and sorting in dorsal interneurons by the Lim-HD proteins Lhx9 and Lhx1. Neural Dev. 2009;4:21.

44. Molander C, Wang HF, Rivero-Melián C, Grant G. Early decline and late restoration of spinal cord binding and transganglionic transport of isolectin B4 from Griffonia simplicifolia I after peripheral nerve transection or crush. Restorative neurology and neuroscience. 1996;10(3):123–33.

45. Zhang Y, Liu CY, Chen WC, Shi YC, Wang CM, Lin S, et al. Regulation of neuropeptide Y in body microenvironments and its potential application in therapies: a review. Cell Biosci. 2021;11(1):151.

46. Li B, Xia M, Harkany T, Verkhratsky A. Long-distance volume transmission for brain-body signaling. Neural Regen Res. 2025.

47. Zubieta JK, Smith YR, Bueller JA, Xu Y, Kilbourn MR, Jewett DM, et al. mu-opioid receptor-mediated antinociceptive responses differ in men and women. J Neurosci. 2002;22(12):5100–7.

48. Pokhilko A, Nash A, Cader MZ. Common transcriptional signatures of neuropathic pain. Pain. 2020;161(7):1542–54.

49. Hastie BA, Riley JL, 3rd, Kaplan L, Herrera DG, Campbell CM, Virtusio K, et al. Ethnicity interacts with the OPRM1 gene in experimental pain sensitivity. Pain. 2012;153(8):1610–9.

50. Staedtler ES, Sapio MR, King DM, Maric D, Ghetti A, Mannes AJ, et al. The mu-opioid receptor differentiates two distinct human nociceptive populations relevant to clinical pain. Cell Rep Med. 2024;5(10):101788.

51. Corder G, Tawfik VL, Wang D, Sypek EI, Low SA, Dickinson JR, et al. Loss of mu opioid receptor signaling in nociceptors, but not microglia, abrogates morphine tolerance without disrupting analgesia. Nat Med. 2017;23(2):164–73.

52. Tsantoulas C, Zhu L, Yip P, Grist J, Michael GJ, McMahon SB. Kv2 dysfunction after peripheral axotomy enhances sensory neuron responsiveness to sustained input. Exp Neurol. 2014;251:115–26.

53. Zhang C, Hu MW, Wang XW, Cui X, Liu J, Huang Q, et al. scRNA-sequencing reveals subtype-specific transcriptomic perturbations in DRG neurons of Pirt(EGFPf) mice in neuropathic pain condition. Elife. 2022;11.

54. Avraham O, Feng R, Ewan EE, Rustenhoven J, Zhao G, Cavalli V. Profiling sensory neuron microenvironment after peripheral and central axon injury reveals key pathways for neural repair. Elife. 2021;10.

55. Feng R, Muraleedharan Saraswathy V, Mokalled MH, Cavalli V. Self-renewing macrophages in dorsal root ganglia contribute to promote nerve regeneration. Proceedings of the National Academy of Sciences of the United States of America. 2023;120(7):e2215906120.

56. Wang K, Wang S, Chen Y, Wu D, Hu X, Lu Y, et al. Single-cell transcriptomic analysis of somatosensory neurons uncovers temporal development of neuropathic pain. Cell Res. 2021;31(8):904–18.

57. Krishnan A, Bhavanam S, Zochodne D. An Intimate Role for Adult Dorsal Root Ganglia Resident Cycling Cells in the Generation of Local Macrophages and Satellite Glial Cells. J Neuropathol Exp Neurol. 2018;77(10):929–41.

58. Mapplebeck JCS, Dalgarno R, Tu Y, Moriarty O, Beggs S, Kwok CHT, et al. Microglial P2X4R-evoked pain hypersensitivity is sexually dimorphic in rats. Pain. 2018;159(9):1752–63.

59. Jancálek R, Dubový P, Svízenská I, Klusáková I. Bilateral changes of TNF-alpha and IL-10 protein in the lumbar and cervical dorsal root ganglia following a unilateral chronic constriction injury of the sciatic nerve. Journal of neuroinflammation. 2010;7:11.

60. Shannon P, Markiel A, Ozier O, Baliga NS, Wang JT, Ramage D, et al. Cytoscape: a software environment for integrated models of biomolecular interaction networks. Genome Res. 2003;13(11):2498–504.

61. Schindelin J, Arganda-Carreras I, Frise E, Kaynig V, Longair M, Pietzsch T, et al. Fiji: an open-source platform for biological-image analysis. Nature methods. 2012;9(7):676–82.

62. Faul F, Erdfelder E, Buchner A, Lang AG. Statistical power analyses using G*Power 3.1: tests for correlation and regression analyses. Behavior research methods. 2009;41(4):1149–60.

63. Zhou Y, Zhou B, Pache L, Chang M, Khodabakhshi AH, Tanaseichuk O, et al. Metascape provides a biologist-oriented resource for the analysis of systems-level datasets. Nature communications. 2019;10(1):1523.

64. Bankhead P, Loughrey MB, Fernandez JA, Dombrowski Y, McArt DG, Dunne PD, et al. QuPath: Open source software for digital pathology image analysis. Sci Rep. 2017;7(1):16878.

65. Chaplan SR, Bach FW, Pogrel JW, Chung JM, Yaksh TL. Quantitative assessment of tactile allodynia in the rat paw. J Neurosci Methods. 1994;53(1):55–63.

66. Gonzalez-Cano R, Boivin B, Bullock D, Cornelissen L, Andrews N, Costigan M. Up-Down Reader: An Open Source Program for Efficiently Processing 50% von Frey Thresholds. Front Pharmacol. 2018;9:433.

67. Martin M. Cutadapt removes adapter sequences from high-throughput sequencing reads. EMBnetjournal. 2011;17(1):3.

68. Dobin A, Davis CA, Schlesinger F, Drenkow J, Zaleski C, Jha S, et al. STAR: ultrafast universal RNA-seq aligner. Bioinformatics. 2013;29(1):15–21.

69. Liao Y, Smyth GK, Shi W. featureCounts: an efficient general purpose program for assigning sequence reads to genomic features. Bioinformatics. 2014;30(7):923–30.

70. Love MI, Huber W, Anders S. Moderated estimation of fold change and dispersion for RNA-seq data with DESeq2. Genome Biol. 2014;15(12):550.

