## supplemental figures and tables for "Sex differences define the molecular and cellular phenotypes of pain resolution in dorsal root ganglia"

### Supplementary Tables

*Supplementary table 1: Ground truth estimation scores of expert annotations*

| Target model | Expert | Dice score | Average dice score | Std dice score | Average std dice score |
|---|---|---|---|---|---|
| ATF3+ neurons | 1 | 0.925 | 0.903 | 0.038 | 0.068 |
| | 2 | 0.940 | | 0.024 | |
| | 3 | 0.843 | | 0.144 | |
| ATF3all neurons | 1 | 0.962 | 0.930 | 0.010 | 0.022 |
| | 2 | 0.931 | | 0.022 | |
| | 3 | 0.897 | | 0.034 | |
| FABP7+ SGCs confocal | 1 | 0.871 | 0.877 | 0.051 | 0.048 |
| | 2 | 0.899 | | 0.040 | |
| | 3 | 0.861 | | 0.053 | |
| IB4+ neurons | 1 | 0.876 | 0.898 | 0.114 | 0.061 |
| | 2 | 0.911 | | 0.031 | |
| | 3 | 0.909 | | 0.038 | |
| NF+ neurons confocal | 1 | 0.966 | 0.971 | 0.012 | 0.011 |
| | 2 | 0.976 | | 0.008 | |
| | 3 | 0.970 | | 0.013 | |

*Supplementary table 2: Performance of models on validation images*

| Model | Model number | Dice score | Average dice score | Uncertainty score | Average uncertainty score |
|---|---|---|---|---|---|
| ATF3+ neurons | 1 | 0.869 | 0.901 | 0.038 | 0.030 |
| | 2 | 0.925 | | 0.026 | |
| | 3 | 0.909 | | 0.027 | |
| ATF3all neurons | 1 | 0.924 | 0.924 | 0.029 | 0.028 |
| | 2 | 0.910 | | 0.028 | |
| | 3 | 0.939 | | 0.026 | |
| FABP7+ SGCs confocal | 1 | 0.857 | 0.849 | 0.046 | 0.051 |
| | 2 | 0.830 | | 0.057 | |
| | 3 | 0.861 | | 0.051 | |
| IB4+ neurons | 1 | 0.887 | 0.859 | 0.026 | 0.040 |
| | 2 | 0.865 | | 0.038 | |
| | 3 | 0.826 | | 0.055 | |
| NF+ neurons confocal | 1 | 0.959 | 0.957 | 0.017 | 0.017 |
| | 2 | 0.950 | | 0.016 | |
| | 3 | 0.961 | | 0.018 | |
| NF+ neurons female | 1 | 0.874 | 0.877 | 0.044 | 0.040 |
| | 2 | 0.920 | | 0.031 | |
| | 3 | 0.836 | | 0.044 | |

*Supplementary table 3: Performance of models on test images*

| Model | Test image | Dice score | Average dice score | Uncertainty score | Average uncertainty score |
|---|---|---|---|---|---|
| ATF3+ neurons | 1 | 0.896 | 0.913 | 0.043 | 0.047 |
| | 2 | 0.912 | | 0.042 | |
| | 3 | 0.929 | | 0.055 | |
| ATF3all neurons | 1 | 0.936 | 0.924 | 0.029 | 0.029 |
| | 2 | 0.905 | | 0.036 | |
| | 3 | 0.932 | | 0.024 | |
| FABP7+ SGCs confocal | 1 | 0.841 | 0.857 | 0.056 | 0.058 |
| | 2 | 0.872 | | 0.059 | |
| | 3 | 0.858 | | 0.060 | |
| IB4+ neurons | 1 | 0.801 | 0.863 | 0.065 | 0.049 |
| | 2 | 0.897 | | 0.043 | |
| | 3 | 0.890 | | 0.040 | |
| NF+ neurons confocal | 1 | 0.935 | 0.964 | 0.025 | 0.016 |
| | 2 | 0.983 | | 0.012 | |
| | 3 | 0.974 | | 0.013 | |
| NF+ neurons female | 1 | 0.888 | 0.904 | 0.044 | 0.043 |
| | 2 | 0.929 | | 0.038 | |
| | 3 | 0.907 | | 0.042 | |
| | 4 | 0.914 | | 0.035 | |
| | 5 | 0.881 | | 0.057 | |

### Supplementary figures

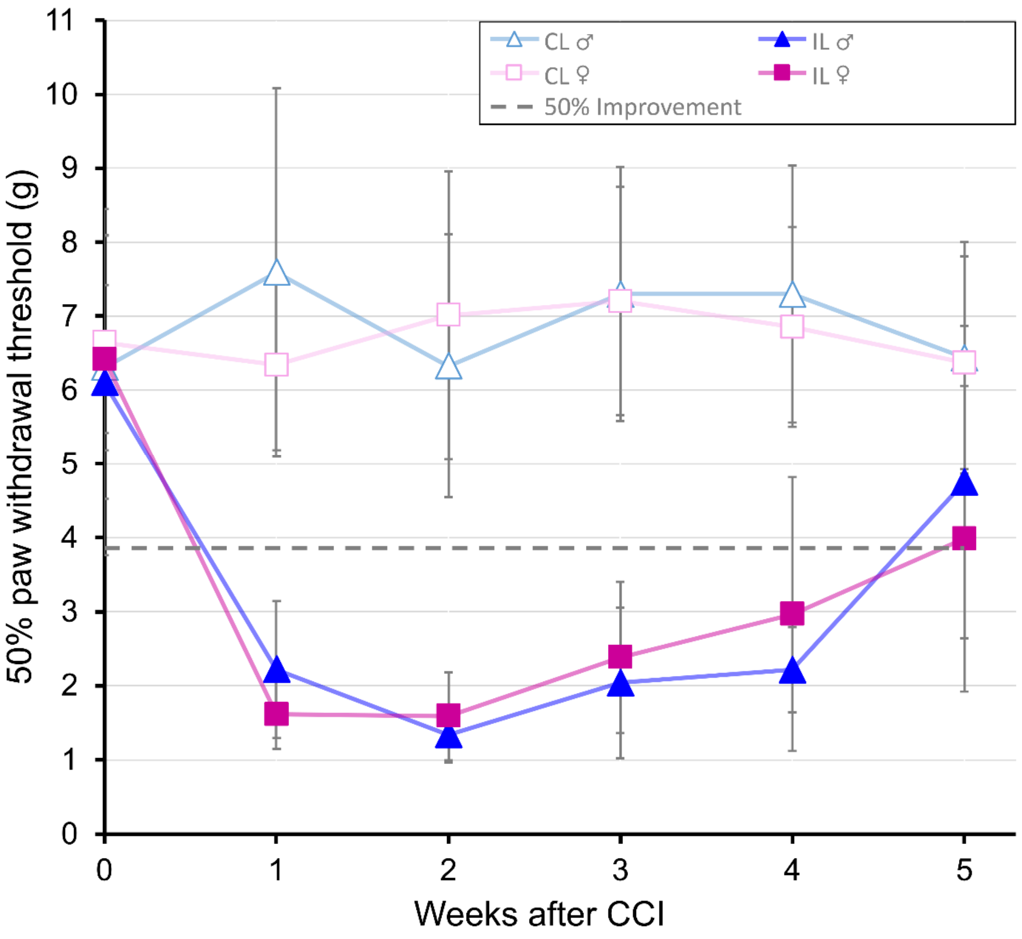

**Figure S1. Mechanical hypersensitivity after CCI.**

Pain resolution was modeled in male and female rats following CCI. The contralateral side served as an internal control. *À priori*, 50% recovery from mechanical hypersensitivity in the von Frey test was defined as the objective time point for ongoing pain resolution. The graph shows paw withdrawal thresholds at baseline and after CCI, with the experimentally determined 50% recovery threshold (grey line) calculated from the peak hypersensitivity value for each sex (blue: males, pink: females; n = 10 males, n = 12 females).

59

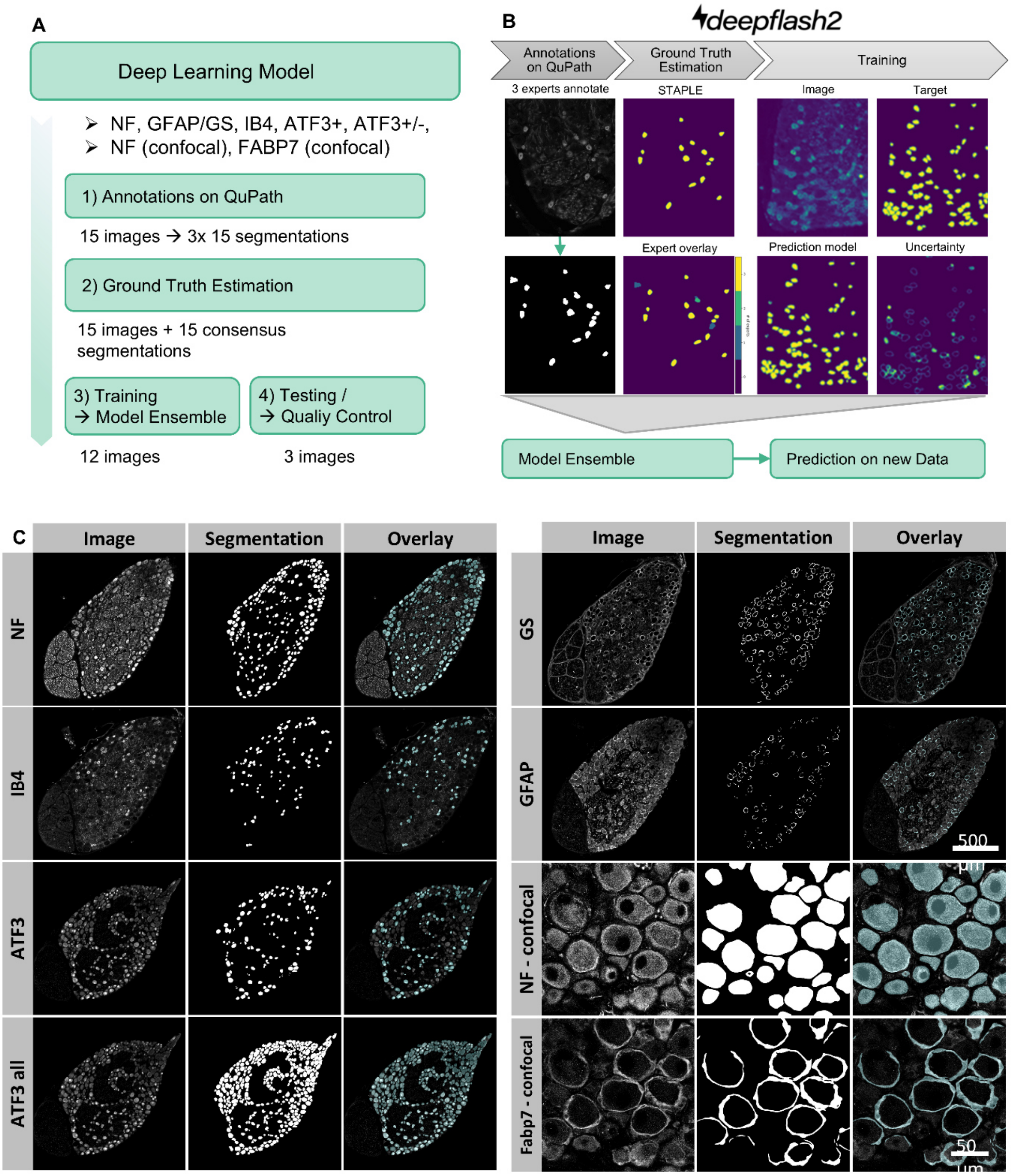

60

61 **Figure S2. Procedure of deep learning-based bioimage segmentation.**

62 (A) Overview of steps in training DL models with deepflash2. (B) Outline of main steps of the deepflash-
63 based image feature segmentation workflow. (C) Examples of bioimages showing immunofluorescence
64 labels, respective feature segmentations, and the corresponding overlay (label in grey, segmentation mask
65 in cyan).

66

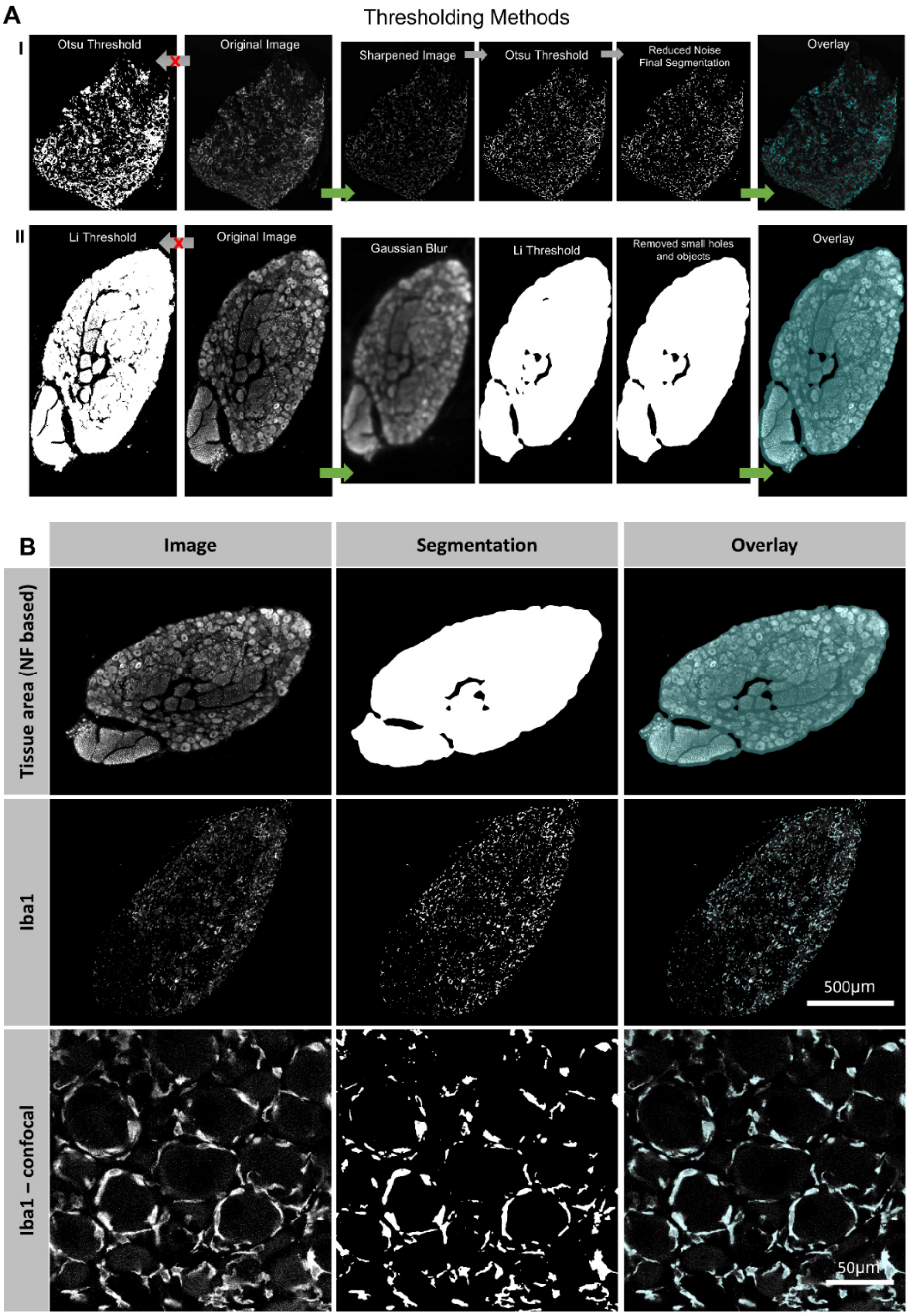

**Figure S3. Representative whole tile- and confocal images showing anti-NF and anti-Iba1 immunofluorescence labelling and respective results from image feature segmentation by advanced thresholding.**

(A) Exemplary images outline computational steps of feature segmentation by advanced thresholding. (B) Exemplary images and their respective segmentations are shown together with an overlay of image (grey) and mask (cyan).

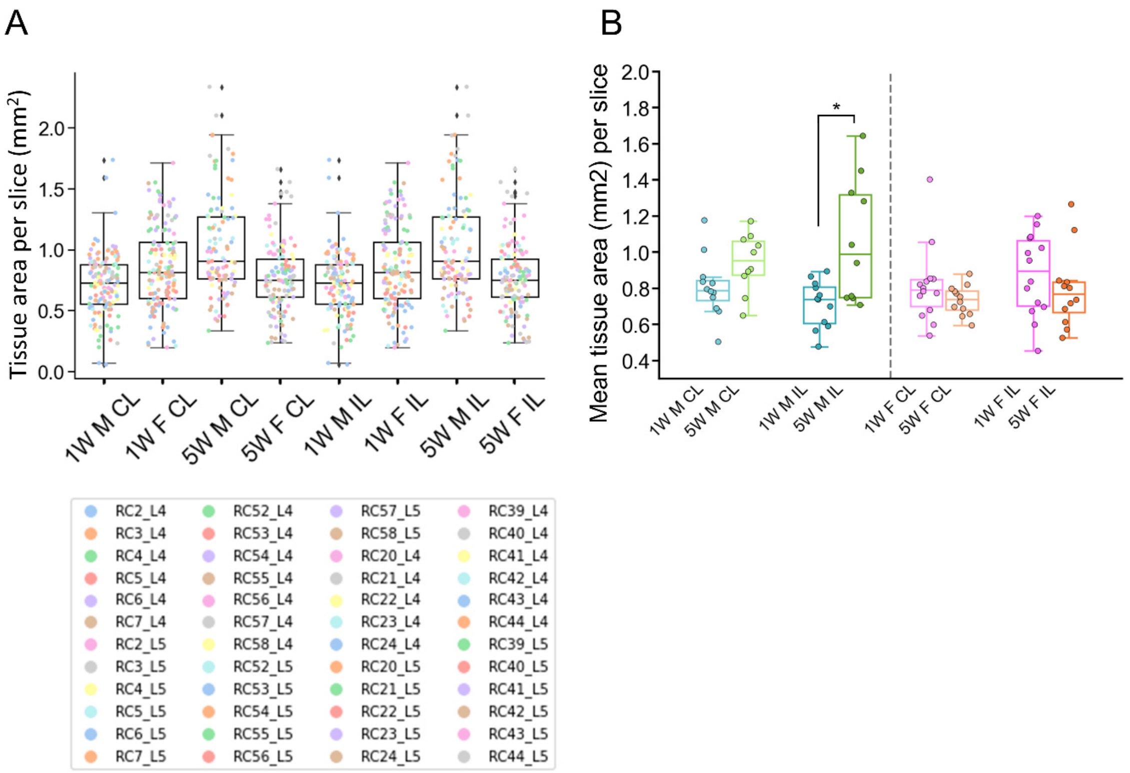

**Figure S4. Documentation of DRG slice parameters.**

(A) Representation of the tissue area for each slice. Each dot represents one slice. (B) Mean tissue area per slice. Data for male (blue/green) or female (pink/orange) rats, 1W/5W after CCI. Each data point represents one DRG. Statistical significance is indicated as *p ≤ 0.05.

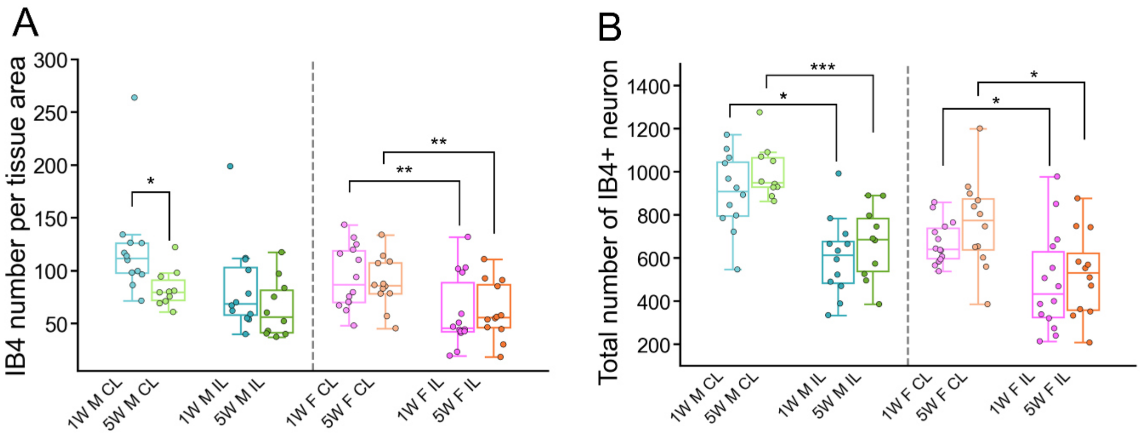

#### Figure S5. IB4 label parameters

Box plots showing (A) the total number of IB4⁺ neurons per tissue area, (B) and total IB4⁺ neuron number as sum of all slices per DRG in male (blue/green) or female (pink/orange) rats, 1 week (1W) or 5 weeks (5W) after CCI. Each data point represents one DRG. Statistical significance is indicated as *p ≤ 0.05; **p ≤ 0.01; ***p ≤ 0.001.

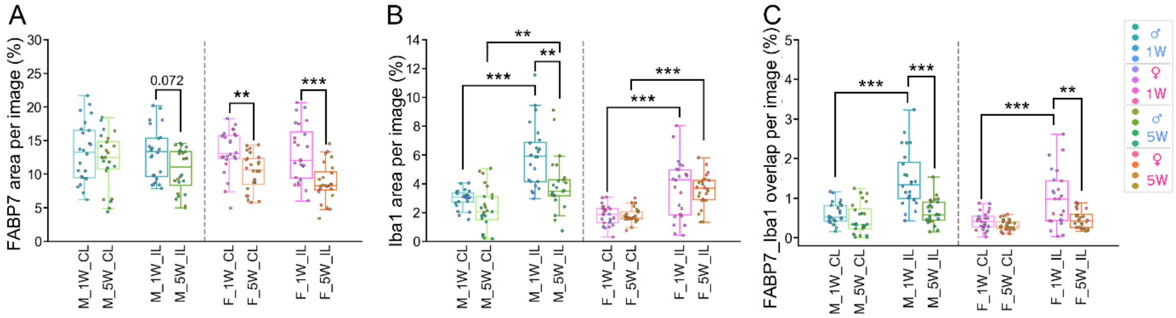

#### Figure S6. Fabp7 and Iba1 label abundance in confocal images.

(A) Percentage of each image that is positive for FABP7, (B) Iba1 or (C) the overlap of both labels in the confocal image dataset. Each dot represents one confocal image. Statistical significance is indicated as *p ≤ 0.05; **p ≤ 0.01; ***p ≤ 0.001. Significance was tested for 1W CL vs. 1W IL, 5W CL vs. 1W IL (“Injury-related changes”), 1W IL vs 5W IL (“Resolution-related changes”), and 1W CL vs. 5W CL (“Lateralization of effects”) for both males and females.

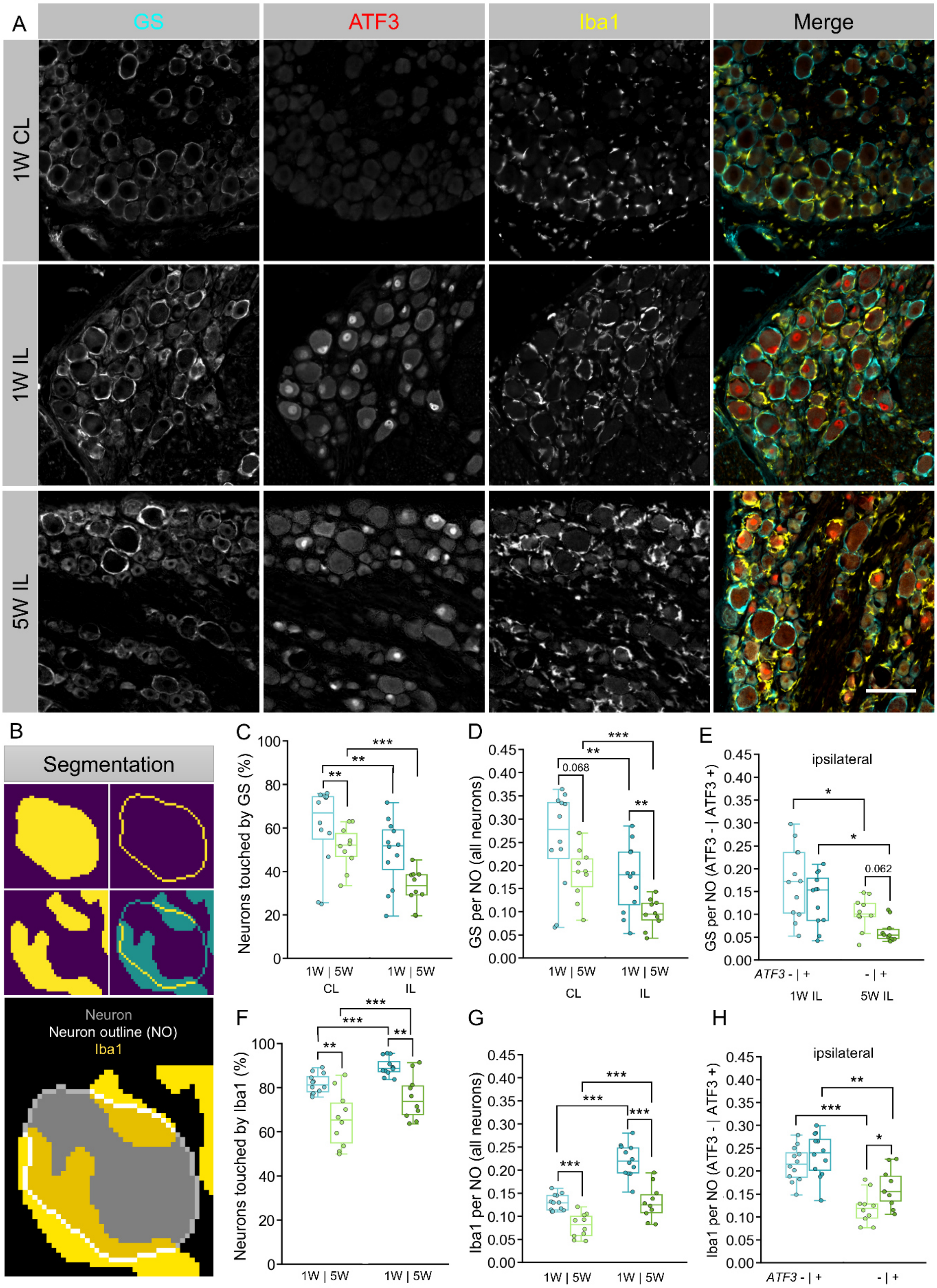
A
GS
ATF3
Iba1
Merge
1W CL
1W IL
5W IL
B
Segmentation
Neuron
Neuron outline (NO)
Iba1
C
Neurons touched by GS (%)
1W | 5W
CL
1W | 5W
IL
D
GS per NO (all neurons)
0.068
E
ipsilateral
GS per NO (ATF3 - | ATF3 +)
0.062
ATF3 - | +
- | +
1W IL
5W IL
F
Neurons touched by Iba1 (%)
G
Iba1 per NO (all neurons)
H
Iba1 per NO (ATF3 - | ATF3 +)

96

**Figure S7. Contact of macrophages and satellite glial cells with the neuron border in ATF3$^+$ and negative neurons.**

(A) Immunofluorescent tile images of rat DRG (L5), representing GS$^+$ SGC (cyan), Iba1$^+$ macrophages (yellow), ATF3$^+$ neurons (red), one or five weeks after CCI. Scale bar: 100µm.

(B) Exemplary images depicting the segmentation of the neuron border (NB) and its overlap with Iba1, as was done for Iba1 and GS.

(C) Percentage of neurons that are touched by GS, where every neuron with any contact is counted as positive. (D) Percentage of the neuron border that overlaps with GS. (E) Separation of the ATF3 positive and negative neuron population of the ipsilateral (IL) side and their percentage of GS contact per NB.

(F) Percentage of neurons that are touched by Iba1, where every neuron with any contact is counted as positive. (G) Percentage of the neuron border that overlaps with Iba1. (H) Separation of the ATF3+ and negative neuron population of the ipsilateral (IL) side and their percentage of macrophage contact (Iba1+) per NB.

(A-G) Each dot represents one DRG. Statistical significance is indicated as *$p \leq 0.05$; **$p \leq 0.01$; ***$p \leq 0.001$. Significance was tested for (C, D, F, G) 1W CL vs. 1W IL (“Injury-related changes”), 1W IL vs 5W IL (“Resolution-related changes”), and 1W CL vs. 5W CL (“Lateralization of effects”) and (E,H) ATF3 negative vs. positive for 1 and 5 weeks (“ATF3 dependency”) and 1W vs. 5W for both ATF3 positive and negative neurons (“Resolution-related changes”) and Bonferroni-corrected per hypothesis.

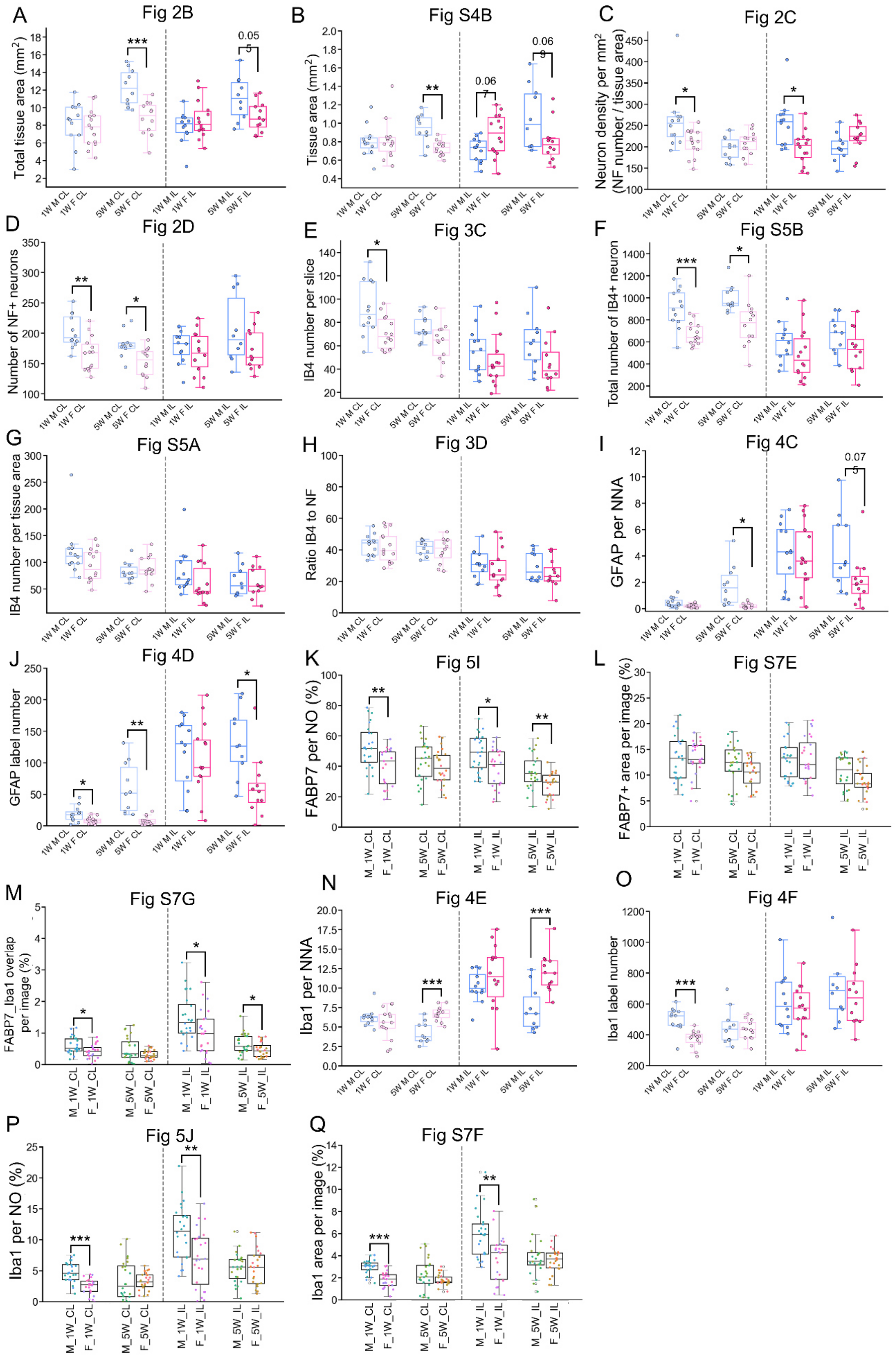
A
Fig 2B
Total tissue area (mm$^2$)
0.05
B
Fig S4B
Tissue area (mm$^2$)
0.06
C
Fig 2C
Neuron density per mm$^2$ (NF number / tissue area)
D
Fig 2D
Number of NF+ neurons
E
Fig 3C
IB4 number per slice
F
Fig S5B
Total number of IB4+ neuron
G
Fig S5A
IB4 number per tissue area
H
Fig 3D
Ratio IB4 to NF
I
Fig 4C
GFAP per NNA
0.07
J
Fig 4D
GFAP label number
K
Fig 5I
FABP7 per NO (%)
L
Fig S7E
FABP7+ area per image (%)
M
Fig S7G
FABP7_Iba1 overlap per image (%)
N
Fig 4E
Iba1 per NNA
O
Fig 4F
Iba1 label number
P
Fig 5J
Iba1 per NO (%)
Q
Fig S7F
Iba1 area per image (%)
1W M CL
1W F CL
5W M CL
5W F CL
1W M IL
1W F IL
5W M IL
5W F IL
M_1W_CL
F_1W_CL
M_5W_CL
F_5W_CL
M_1W_IL
F_1W_IL
M_5W_IL
F_5W_IL

**Figure S8. Female vs. male comparisons.**

Shown are boxplots as indicated, rearranged to highlight sex-specific comparisons. Significance is indicated as $*p \leq 0.05$; $**p \leq 0.01$; $***p \leq 0.001$.

M = male; F = female; W = weeks; IL = ipsilateral; CL = contralateral; NNA = neuron-near area; NO = neuron segmentation outline. Significance was tested as male vs. female for each timepoint on the contralateral side ("baseline differences") and on the ipsilateral side ("injury effect"). Bonferroni correction for each hypothesis.

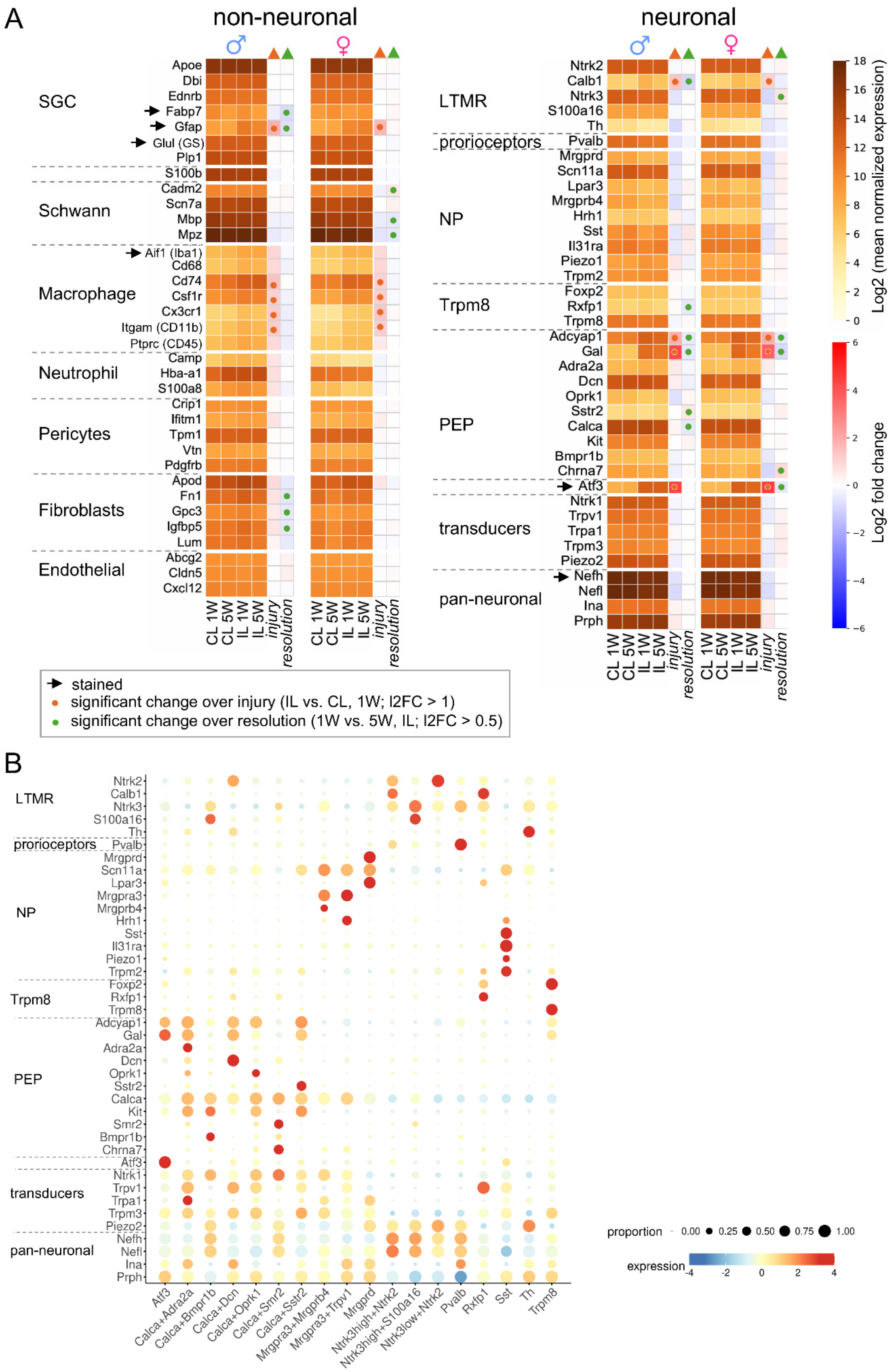
A
non-neuronal
neuronal
SGC
Apoe
Dbi
Ednrb
Fabp7
Gfap
Glul (GS)
Plp1
S100b
Schwann
Cadm2
Scn7a
Mbp
Mpz
Macrophage
Aif1 (Iba1)
Cd68
Cd74
Csf1r
Cx3cr1
Itgam (CD11b)
Ptprc (CD45)
Neutrophil
Camp
Hba-a1
S100a8
Pericytes
Crip1
Ifitm1
Tpm1
Vtn
Pdgfrb
Fibroblasts
Apod
Fn1
Gpc3
Igfbp5
Lum
Endothelial
Abcg2
Cldn5
Cxcl12
CL 1W
CL 5W
IL 1W
IL 5W
injury
resolution
LTMR
Ntrk2
Calb1
Ntrk3
S100a16
Th
proprioceptors
Pvalb
NP
Mrgprd
Scn11a
Lpar3
Mrgprb4
Hrh1
Sst
Il31ra
Piezo1
Trpm2
Trpm8
Foxp2
Rxfp1
Trpm8
PEP
Adcyap1
Gal
Adra2a
Dcn
Oprk1
Sstr2
Calca
Kit
Bmpr1b
Chrna7
Atf3
transducers
Ntrk1
Trpv1
Trpa1
Trpm3
Piezo2
pan-neuronal
Nefh
Nefl
Ina
Prph
Log2 (mean normalized expression)
Log2 fold change
stained
significant change over injury (IL vs. CL, 1W; l2FC > 1)
significant change over resolution (1W vs. 5W, IL; l2FC > 0.5)
B
LTMR
Ntrk2
Calb1
Ntrk3
S100a16
Th
proprioceptors
Pvalb
NP
Mrgprd
Scn11a
Lpar3
Mrgpra3
Mrgprb4
Hrh1
Sst
Il31ra
Piezo1
Trpm2
Trpm8
Foxp2
Rxfp1
Trpm8
PEP
Adcyap1
Gal
Adra2a
Dcn
Oprk1
Sstr2
Calca
Kit
Smr2
Bmpr1b
Chrna7
Atf3
transducers
Ntrk1
Trpv1
Trpa1
Trpm3
Piezo2
pan-neuronal
Nefh
Nefl
Ina
Prph
Atf3
Calca+Adra2a
Calca+Bmpr1b
Calca+Dcn
Calca+Oprk1
Calca+Smr2
Calca+Sstr2
Mrgpra3+Mrgprb4
Mrgpra3+Trpv1
Mrgprd
Ntrk3high+Ntrk2
Ntrk3high+S100a16
Ntrk3low+Ntrk2
Pvalb
Rxfp1
Sst
Th
Trpm8
proportion
0.00
0.25
0.50
0.75
1.00
expression
-4
-2
0
2
4

**Figure S9. Cell type marker expression.**

(A) Heatmaps of selected cell type markers for non-neuronal and neuronal cells expressed in male and female rats for each condition. (B) Expression of each neuronal marker in their subgroup is shown according to categories described by the harmonized cross-species cell atlas of dorsal root ganglia (Bhuiyan et al., 2024).

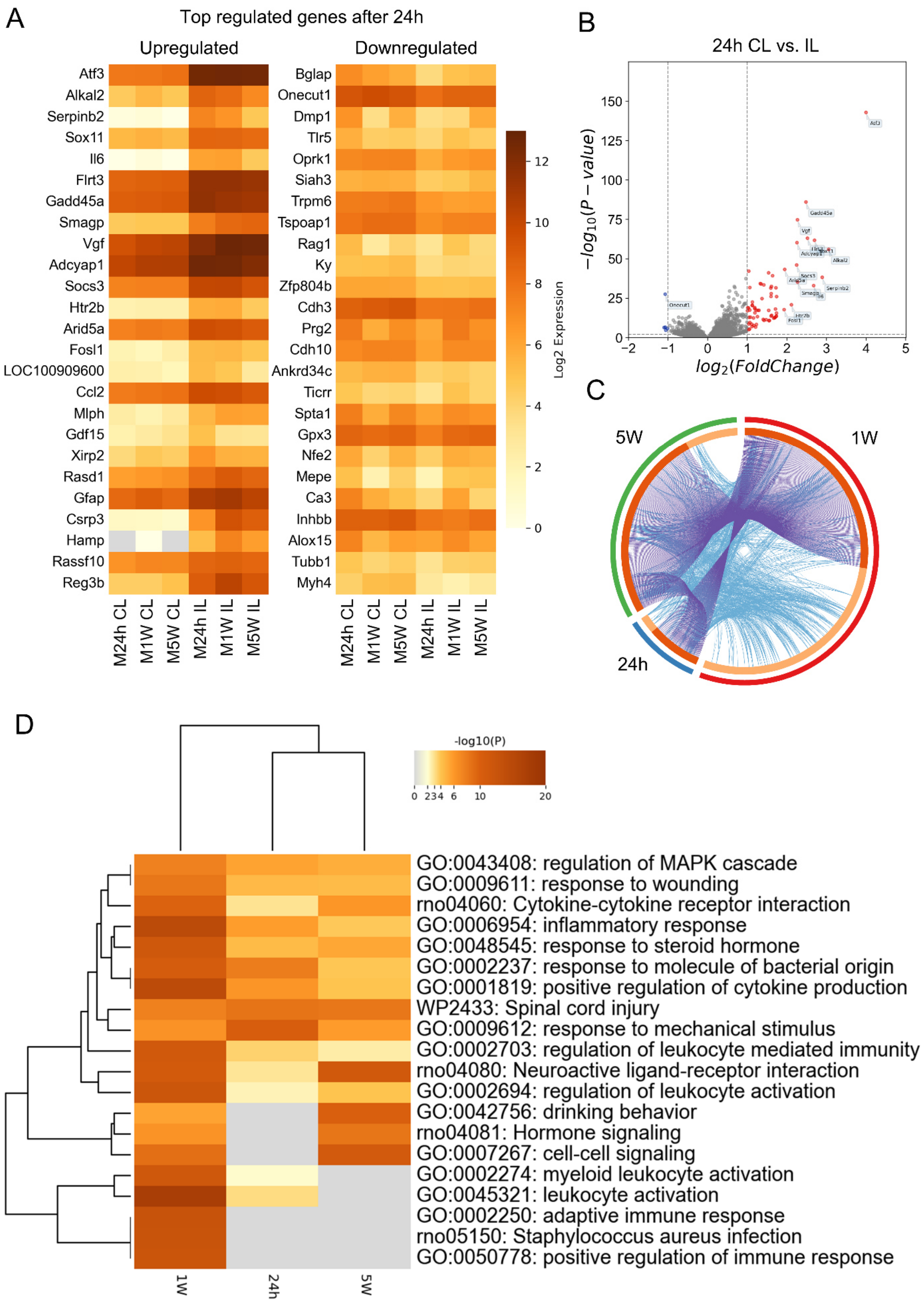
A
Top regulated genes after 24h
Upregulated
Downregulated
Atf3
Alkal2
Serpinb2
Sox11
Il6
Flrt3
Gadd45a
Smagp
Vgf
Adcyap1
Socs3
Htr2b
Arid5a
Fosl1
LOC100909600
Ccl2
Mlph
Gdf15
Xirp2
Rasd1
Gfap
Csrp3
Hamp
Rassf10
Reg3b
Bglap
Onecut1
Dmp1
Tlr5
Oprk1
Siah3
Trpm6
Tspoap1
Rag1
Ky
Zfp804b
Cdh3
Prg2
Cdh10
Ankrd34c
Ticrr
Spta1
Gpx3
Nfe2
Mepe
Ca3
Inhbb
Alox15
Tubb1
Myh4
M24h CL
M1W CL
M5W CL
M24h IL
M1W IL
M5W IL
Log2 Expression
B
24h CL vs. IL
$-log_{10}(P-value)$
$log_2(FoldChange)$
C
5W
1W
24h
D
-log10(P)
GO:0043408: regulation of MAPK cascade
GO:0009611: response to wounding
rno04060: Cytokine-cytokine receptor interaction
GO:0006954: inflammatory response
GO:0048545: response to steroid hormone
GO:0002237: response to molecule of bacterial origin
GO:0001819: positive regulation of cytokine production
WP2433: Spinal cord injury
GO:0009612: response to mechanical stimulus
GO:0002703: regulation of leukocyte mediated immunity
rno04080: Neuroactive ligand-receptor interaction
GO:0002694: regulation of leukocyte activation
GO:0042756: drinking behavior
rno04081: Hormone signaling
GO:0007267: cell-cell signaling
GO:0002274: myeloid leukocyte activation
GO:0045321: leukocyte activation
GO:0002250: adaptive immune response
rno05150: Staphylococcus aureus infection
GO:0050778: positive regulation of immune response
1W
24h
5W

135

**Figure S10. Transcriptomic changes 24h after CCI.**

All figures in this list are based on comparing transcriptome data from the ipsi- with the contralateral side, at 24h, 1 or 5 weeks after CCI in males.

(A) Heatmaps illustrating the top upregulated (left panel) and downregulated (right panel) genes in males at 24h post injury.

(B) Volcano plot highlighting the most significantly upregulated (red) and downregulated (blue) genes. Genes with an absolute log2fold change ≥ 1 and an adjusted p-value <0.01 were considered significantly differentially expressed.

(C) Circos plot (https://circos.ca/) of overlaps between gene lists of significant differential regulation of gene expression, 24h, 1W and 5W after CCI. Purple lines link identical genes between conditions and blue lines link genes with shared ontology terms. Created with metascape.org.

(D) Heatmap of enriched terms in pathway analysis coded by p-value, created with metascape.org.

CL, contralateral; Il, ipsilateral; W = weeks; M = male.

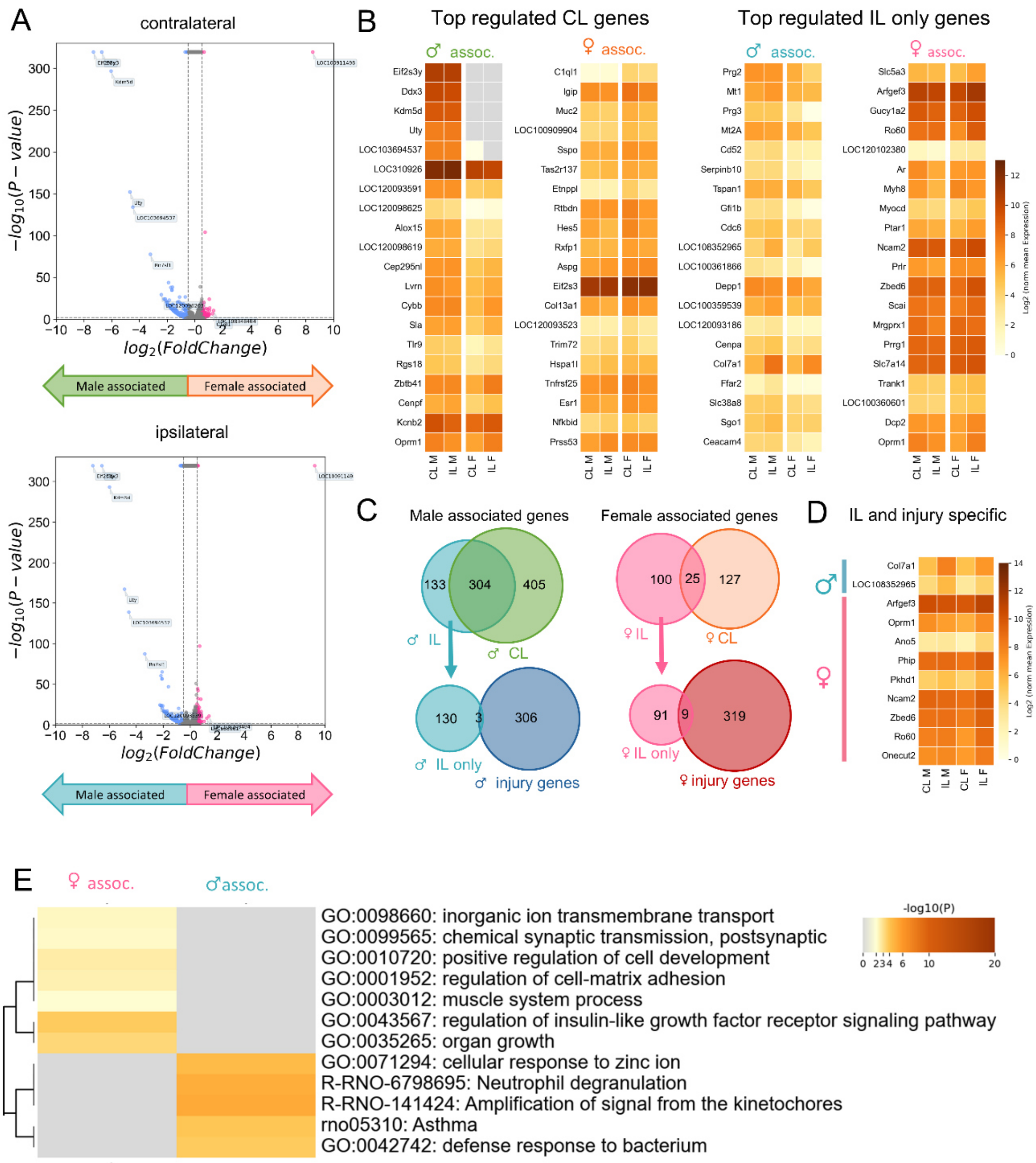

**Figure S11. Sex differences in gene expression at 5 weeks post CCI.**
(A) Volcano plots showing genes that are more strongly expressed and associated with males or females, comparing the contra- (CL) or ipsilateral (IL) side, 5 weeks after CCI. (B) Heatmap of the top male- and female-associated regulated genes on the contralateral side or, ones exclusively occurring on the ipsilateral side, respectively. (C) Venn diagram showing the overlap of male and female-associated genes on the contra- and ipsilateral sides. To find injury-specific differences, we overlapped genes exclusively occurring sex-associated on the ipsilateral side with genes that are also differentially regulated between the ipsi- and contralateral side (injury genes) in each sex. (D) Heatmap of the resulting injury and IL-specific genes that are differentially expressed between both sexes. (E) Pathway analysis of male- or female-associated, IL-specific genes using https://metascape.org.

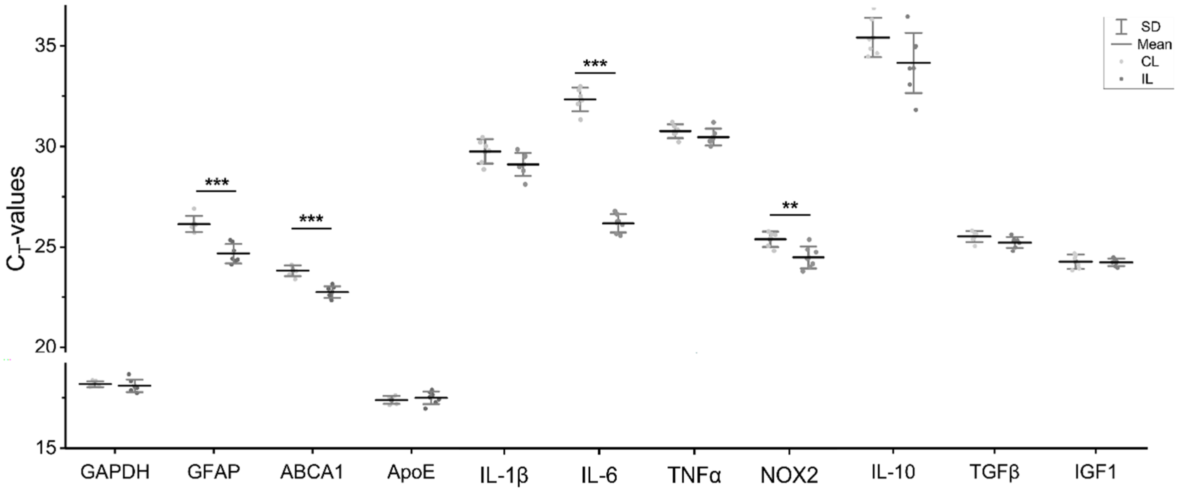

**Figure S12. qPCR of selected marker genes for RNA control.**

Means of the raw CT-values per target gene on the ipsi- and contralateral sides of L5 DRGs from the male cohort one week after CCI. The data was normally distributed, and significance was tested with a two-tailed t-test. Significance is indicated as follows: *p ≤0.05; **p ≤0.01; ***p ≤0.001.

**Reference**

Bhuiyan, S.A., M. Xu, L. Yang, E. Semizoglou, P. Bhatia, K.I. Pantaleo, I. Tochitsky, A. Jain, B. Erdogan, S. Blair, V. Cat, J.M. Mwirigi, I. Sankaranarayanan, D. Tavares-Ferreira, U. Green, L.A. McIlvried, B.A. Copits, Z. Bertels, J.S. Del Rosario, A.J. Widman, R.A. Slivicki, J. Yi, R. Sharif-Naeini, C.J. Woolf, J.K. Lennerz, J.L. Whited, T.J. Price, R.W. Gereau, and W. Renthal. 2024. Harmonized cross-species cell atlases of trigeminal and dorsal root ganglia. *Science Advances*. 10:eadj9173.
